# Symmetry facilitated the evolution of heterospecificity and high-order stoichiometry in vertebrate hemoglobin

**DOI:** 10.1101/2024.07.24.604985

**Authors:** Carlos R. Cortez-Romero, Jixing Lyu, Arvind S. Pillai, Arthur Laganowsky, Joseph W. Thornton

**Affiliations:** Department of Cell and Molecular Biology; Department of Ecology and Evolution; Department of Human Genetics, University of Chicago, Chicago, IL, 60637.; Department of Chemistry, Texas A&M University, College Station, TX, 77843.; Department of Institute of Protein Design, University of Washington, Seattle, WA, 98195

**Keywords:** protein evolution, molecular complexes, multimeric proteins, heterodimers, evolution of specificity, gene duplication, isology

## Abstract

Many proteins form paralogous multimers – molecular complexes in which evolutionarily related proteins are arranged into specific quaternary structures. Little is known about the mechanisms by which they acquired their stoichiometry (the number of total subunits in the complex) and heterospecificity (the preference of subunits for their paralogs rather than other copies of the same protein). Here we use ancestral protein reconstruction and biochemical experiments to study historical increases in stoichiometry and specificity during the evolution of vertebrate hemoglobin (Hb), a α_2_β_2_ heterotetramer that evolved from a homodimeric ancestor after a gene duplication. We show that the mechanisms for this evolutionary transition were simple. One hydrophobic substitution in subunit β after the gene duplication was sufficient to cause the ancestral dimer to homotetramerize with high affinity across a new interface. During this same interval, a single-residue deletion in subunit α at the older interface conferred specificity for the heterotetrameric form and the *trans*-orientation of subunits within it. These sudden transitions in stoichiometry and specificity were possible because the interfaces in Hb are isologous – involving the same surface patch on interacting subunits, rotated 180° relative to each other. This architecture amplifies the impacts of individual mutations on stoichiometry and specificity, especially in higher-order complexes, and allows single substitutions to differentially affect heteromeric vs homomeric interactions. Our findings suggest that elaborate and specific symmetrical molecular complexes may often evolve via simple genetic and physical mechanisms.

**Significance statement:** Many molecular complexes are made up of proteins related by gene duplication, but how these assemblies evolve is poorly understood. Using ancestral protein reconstruction and biochemical experiments, we dissected how vertebrate hemoglobin, which comprises two copies each of two related proteins, acquired this architecture from a homodimeric ancestor. Each aspect of this transition – from dimer to tetramer and homomer to heteromer – had a simple genetic basis: a single-site amino acid change in each protein drove these changes in size and specificity. These transitions were possible because hemoglobin’s architecture is symmetric, which amplified the effect of small biochemical changes on the assembly of the complex. Many protein complexes are symmetrical, suggesting that they too may have evolved via simple genetic mechanisms.

**Classification: Major classification** - Biological Sciences **Minor classifications** – Biochemistry/Evolution

## INTRODUCTION

Protein multimers – associations of multiple protein subunits arranged in specific quaternary architectures – carry out most biochemical functions in living cells (1, 2). The mechanisms by which these complexes evolved their stoichiometry and specificity present some puzzling questions (2–10). Multimers assemble via interfaces that typically contain dozens of sterically and electrostatically complementary residues, and higher-than-dimeric stoichiometries (tetramers, octamers, etc.) use several such interfaces on each subunit (11). This seems to imply that many sequence substitutions would be required for a new multimeric assembly to originate during evolution.

A second complication is that many multimers are composed of paralogs -- proteins related to each other by gene duplication (12). Paralogs are genetically and structurally indistinguishable when generated by duplication, so initially they assemble indiscriminately into homomers and heteromers. Most complexes, however, have evolved specificity for either the homomeric or heteromeric form, with the latter being the most common outcome (12, 13). How specificity evolves is unclear, because mutations that affect multimerization are expected to cause correlated effects on the affinities of homomerization and heteromerization (6,12,14). The structural similarity of paralogs seems to imply that substitutions in both paralogs are required to confer any specificity at all. This complication is magnified for higher-order paralogous multimers, in which one might expect that every interface must evolve specificity to mediate assembly into the complex’s particular architecture.

A critical factor in the evolution of specificity and high-order stoichiometry may be whether a multimer assembles through symmetrical interfaces. In many complexes, identical or paralogous subunits bind each other using an isologous interface – a form of symmetry in which a surface patch on one subunit binds to the same patch on its partner but rotated 180 degrees relative to each other (1). Isologous complexes might, in principle, have the potential to evolve changes in stoichiometry and specificity through simpler mechanisms than nonisologous head-to-tail interfaces. A single substitution appears twice across the interface(s) of an isologous homodimer or heterotetramer, four times in a homotetramer, etc. (Fig. 1A). Mutations that weakly affect affinity on their own can therefore confer large effects on the assembly of isologous multimers (1,5,9,15–17). Isology also changes the way that mutations can affect specificity. In a nonisologous interface, specificity requires mutations on both surfaces so that the tails are recognizably different from each other and each head prefers one tail over the other. In an isologous interface, however, a substitution on the surface of just one subunit has the potential to differentially affect the affinity of each kind of complex, because it will appear twice in the interface of a homomer, once in the heteromer, and not at all in the other homomer (Fig. 1A).

**Figure 1.**
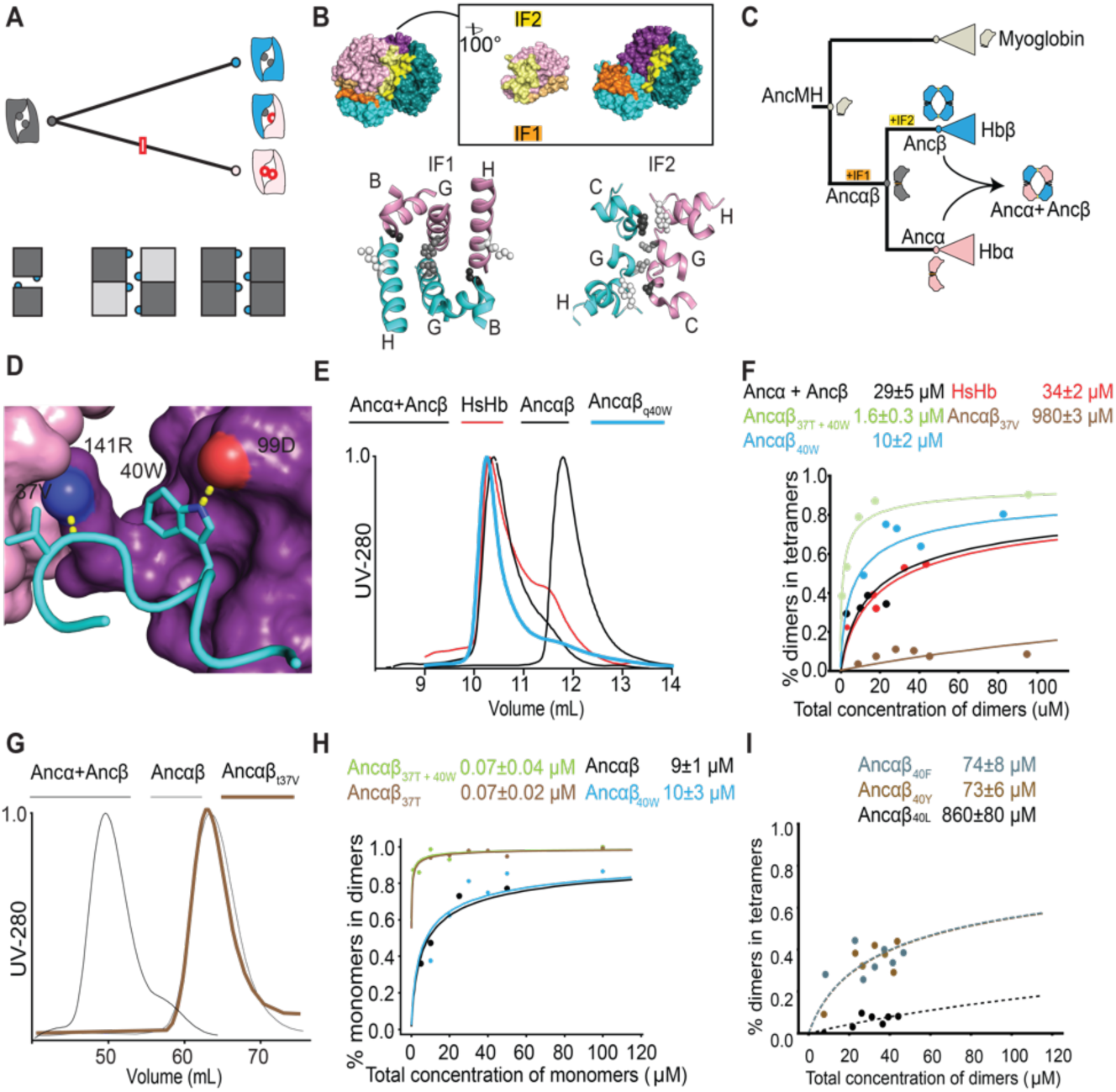
A single substitution confers tetramerization on an ancestral dimer. **(A)** A substitution in one subunit can potentially affect specificity and stoichiometry in an isologous interface. *Top*: After duplication of an isologous homodimer (gray), a substitution that occurs in one paralog (red box) appears twice in the interface of a homodimer (red circles), once in a heterodimer, and not at all in the other homodimer (blue). *Bottom*: One substitution (blue circle) in an isologous interface appears twice in a homodimer (*left*), twice in a heterotetramer (*middle*), and four times in a homotetramer (*right*), multiplying its effects on affinity. Dark and light gray, paralogous subunits. **(B)** *Top*: Interfaces in the human Hb heterotetramer (PDB 4HHB). Pink, Hbα; blue, Hbβ; α_1_ and β_1_ are in lighter hues than α_2_ and β_2_. IF1 surfaces (orange) mediate α_1_-β_1_ and α_2_-β_2_ interactions; yellow surfaces (IF2) mediate α_1_-β_2_ and α_2_-β_1_ interactions. Only interfaces involving α_1_ are shown. Inset, α_1_ subunit rotated away from the rest of the tetramer to show IF1 and IF2. *Bottom*: Isology of IF1 and IF2. Helices contributing to each interface are shown and labeled. Balls and sticks: on each helix, one residue’s side chain is shown to visualize symmetry. **(C)** Evolution of tetrameric stoichiometry on the phylogeny of Hb and related globins. Icons, oligomeric states determined by experimental characterization of reconstructed ancestral proteins (16). Acquisition of interfaces of IF1 and IF2 is shown (16). **(D)** Key residues V37 and W40 that were substituted in Ancb. Cyan cartoon helix, b_1_ subunit. Pink and violet surfaces, a subunits that interact with b_1_ via IF1 and IF2, respectively. Dotted lines to red or blue spheres, hydrogen bonds to oxygen or nitrogen atoms, respectively (PDB 4HHB). **(E,G)** Effect of historical substitutions on stoichiometry, as measured by size exclusion chromatography. The ancestral dimer Ancαβ and the tetramers Ancα+Ancβ and human hemoglobin (HsHb) are shown for comparison. Protein concentration at 100 mM (E) or 1 mM (G). **(F)** Dimer-to-tetramer affinity of reconstructed ancestral Hb subunits containing historical substitutions q40W and t37V, measured by native mass spectrometry across a titration series. Points, fraction of dimers that are incorporated into tetramers. Lines, best-fit binding curves. Estimated Kd and 95% confidence interval are shown. **(H)** Effect of historical substitutions on monomer-dimer affinity measured by native MS. **(I)** Effect on dimer-tetramer affinity of nonhistorical hydrophobic mutations in at residue 40, measured by native MS.

Little is known about the historical evolution of heterospecific complexes or the role of symmetry in this process, especially in high-order complexes. Biochemical and protein engineering studies have addressed the determinants of binding affinity in both homomeric and heteromeric interfaces of extant proteins (20–26). But the genetic and structural mechanisms by which those interactions were acquired long ago are often different from their derived forms in the present (27). Ancestral sequence reconstruction (ASR) can address this limitation by experimentally characterizing the effects of historical sequence changes when introduced into ancestral proteins. ASR has been used to understand the evolution of specificity after duplication in head-to-tail paralogous heteromers (18,19) and in multimers composed of unrelated proteins, which are by definition asymmetrical (26). But we know of no studies that have addressed how isologous heteromers historically evolved their specificity or how specificity in high-order complexes was acquired. A recent in silico analysis predicted that it should be possible for specificity in heterodimers to evolve rapidly after gene duplication through small perturbations in binding energy (28), but the underlying mechanisms and historical relevance of this phenomenon are unknown.

Here we use ASR to study the evolution of higher-order stoichiometry and specificity in vertebrate hemoglobin (Hb), the major carrier of oxygen in the blood of jawed vertebrates. Hb is a paralogous α_2_β_2_ heterotetramer ((17), Fig. 1B), assembly of which is mediated by two distinct and isologous interface patches (IF1 and IF2). Each subunit of the tetramer uses its IF1 to bind IF1 of a paralogous subunit; two of these heterodimers bind to each other using the IF2 on each subunit to make the tetramer ((29), Fig. 1B). Hb⍺ and Hbβ descend from a gene duplication deep in the vertebrate lineage (Fig. 1C), and their sequences retain sufficient phylogenetic signal to allow high-confidence reconstruction of ancestral Hb protein sequences. Using ASR, we recently showed experimentally that extant Hb evolved its heterotetrameric architecture in two phases from a monomeric precursor via a homodimeric intermediate (17). In the first phase, prior to the gene duplication that yielded paralogous ⍺ and β lineages, a monomeric ancestor evolved the capacity to homodimerize with moderate affinity across IF1. In the second phase – after the gene duplication but before the last common ancestor of all vertebrates – binding across IF2 was acquired, yielding the tetrameric stoichiometry, and specificity for the heteromeric form α_2_β_2_ also evolved (Fig. 1C).

Here we characterize the genetic and physical mechanisms that mediated the evolutionary transition from homodimer to heterotetramer in this second phase. By experimentally characterizing reconstructed ancestral hemoglobin subunits and the effects of historical sequence changes on them, we address the following questions: 1) How many substitutions were required to confer tetrameterization across IF2, and what thermodynamic and structural mechanisms mediated their effects? 2) Did the evolution of specificity for the heterotetrameric form require sequence changes at one or both interfaces, in one or both subunits, and what physical mechanisms drove the acquisition of this specificity? 3) How did the symmetry of Hb’s two interfaces affect this evolutionary transition to a high-order, heterospecific architecture? 4) Does a mutational propensity favor increased molecular complexity during the evolution of isologous complexes?

## RESULTS

### Evolution of tetrameric stoichiometry

We first sought to identify the historical sequence changes that conferred tetramerization after duplication of the ancestral homodimer Ancαβ. We focused on the branch leading from the duplication of Ancαβ to Ancβ (the Hbβ subunit in the last common ancestor of jawed vertebrates), because Ancβ heterotetramerizes with Ancα (the Hbα subunit in the jawed vertebrate ancestor) and, like extant Hbβs, also homotetramerizes with itself. We previously found two amino acid replacements that occurred on the branch which, if introduced together into Ancαβ, are sufficient to confer high-affinity assembly into homotetramers (17). One of these (q40W) is buried in the IF2 interface, whereas the other (t37V) makes contacts across both IF1 and IF2 (Fig. 1D. 4, using lower and upper case to denote ancestral and derived amino acids, respectively). W40 is strictly conserved in Hbβ subunits throughout the jawed vertebrates, and V37 is conserved in Hbβ of most taxa (**Fig. S1**).

Here we isolated the individual contributions of each amino acid changer by introducing them singly into Ancαβ and characterizing their effect on assembly into tetramers using size-exclusion chromatography (SEC) and native mass spectrometry (nMS) (30,31). We found that q40W alone is sufficient to recapitulate the evolution of Hb’s tetrameric stoichiometry. Ancαβ forms only dimers in SEC at 100 µM of total protein subunits; by contrast, the mutant Ancαβ_q40W_ is tetrameric, with occupancy of the tetramer similar to that observed in the derived Ancα + Ancβ complex and human Hb (Fig. 1E). We used nMS across a titration series to measure the affinity with which dimers associate into tetramers and found that the tetramerization affinity of Ancαβ_q40W_ (Kd 10 µM) is stronger than that of Ancα + Ancβ (29 µM) and human Hb (34 µM) (Fig. 1F). The conclusion that q40W is sufficient to confer tetramerization is robust to statistical uncertainty about the ancestral sequence: similar experiments using a different reconstruction of Ancαβ that incorporates alternative residues at all ambiguously reconstructed sites yields almost identical results (Fig. S2).

The other historical replacement, t37V, is not sufficient to confer tetramerization. Mutant Ancαβ_t37V_ confers no detectable tetramer occupancy by SEC, even at 1 mM (Fig. 1G), and it displays no measurable affinity to form tetramers using nMS (Fig. 1H). When combined with q40W, however, t37V does increase affinity of the dimer-tetramer transition by a factor of 6 compared to the effect of q40W alone (Fig. 1D; Fig. S3).

In principle, a sequence change could also facilitate tetramerization by increasing affinity of the monomer-to-dimer transition; by increasing the effective concentration of dimers, more tetramers would be produced at a given protein concentration, even if affinity of the dimer-tetramer transition were unchanged. Using nMS, we found that t37V improves the monomer-dimer affinity of Ancαβ by >100-fold (Fig. 1H; Fig. S3). Substitution q40W, in contrast, has no effect on monomer-dimer affinity. These findings are consistent with the structural location of these residues -- t37V contributes to both IF1 and IF2 and q40W to IF2 only -- and they explain why t37V does not confer tetramerization on its own but enhances the impact of q40W.

A likely physical mechanism for the effect of q40W is that tryptophan’s bulky hydrophobic side chain nestles into a hydrophobic divot on the IF2 surface of the facing subunit, and is further strengthened by a hydrogen bond to 102D (32). To test this hypothesis, we identified alternative amino acid replacements with similar biochemical properties and measured whether they also could have caused Ancαβ to evolve into a tetramer. Like tryptophan, the bulky hydrophobic residues phenylalanine or tyrosine at this position also confer tetramerization, albeit at affinity slightly weaker than q40W but similar to that of Ancα+Ancβ and human Hb. The greater affinity of tryptophan may be due to its longer side chain, which buries more hydrophobic surface area across the interface; the hydrogen bond with 102D could make a small contribution but is not necessary, because phenylalanine confers tetramerization but provides no hydrogen bond donor. Leucine, in contrast, which has a smaller volume and no hydrogen bonding capacity, confers no measurable tetramerization (Fig. 1I). High-affinity homotetramerization could therefore have evolved via any of three different aromatic replacements at site 40.

Taken together, these data indicate that replacing the amino acid at a single residue position was sufficient to confer tetramerization during historical Hb evolution, and several alternative replacements at the same site could also have caused the acquisition of this higher-order stoichiometry.

### Isology facilitated IF2 evolution

How could a single amino acid replacement cause such a dramatic change in stoichiometry? The Hb tetramer can be viewed as two heterodimers, each of which is mediated by isologous assembly across IF1 (the larger interface); these heterodimers then bind to each other isologously across IF2. We hypothesized that this doubly symmetrical architecture allowed substitution q40W to confer the dimer-tetramer evolutionary transition, because isology causes the derived amino acid to appear four times in the homotetramer and twice in the heterotetramer.

If this hypothesis is correct, then assembly across IF2 by the derived Hb protein should require assembly across IF1 to multiply the intrinsic affinity of IF2 (Fig. 1A). We tested this prediction by introducing q40W into Ancαβ but doing so under conditions that prevent assembly across IF1. We first compromised dimerization across IF1 genetically by reverting the IF1 surface to the ancestral states of the monomeric ancestor AncMH; these mutations abolish dimer occupancy, leaving a monomers-only population at 20 µM (Fig. 2A). We then introduced q40W into these IF1-ablated mutants and assessed stoichiometry using nMS. As predicted, these proteins do not form any observable dimers or tetramers (detection limit ∼1 µM) (Fig. S4). Similar results are found when we used the combination of t37V/q40W to confer association across IF2 or the mutation P127R – which introduces unsatisfied positive charges into IF1 -- to compromise IF1. (Fig. 2B). The IF2 mutations do not compromise heme binding or solubility, because the mutant proteins are purifiable and heme-bound in nMS.

**Figure 2.**
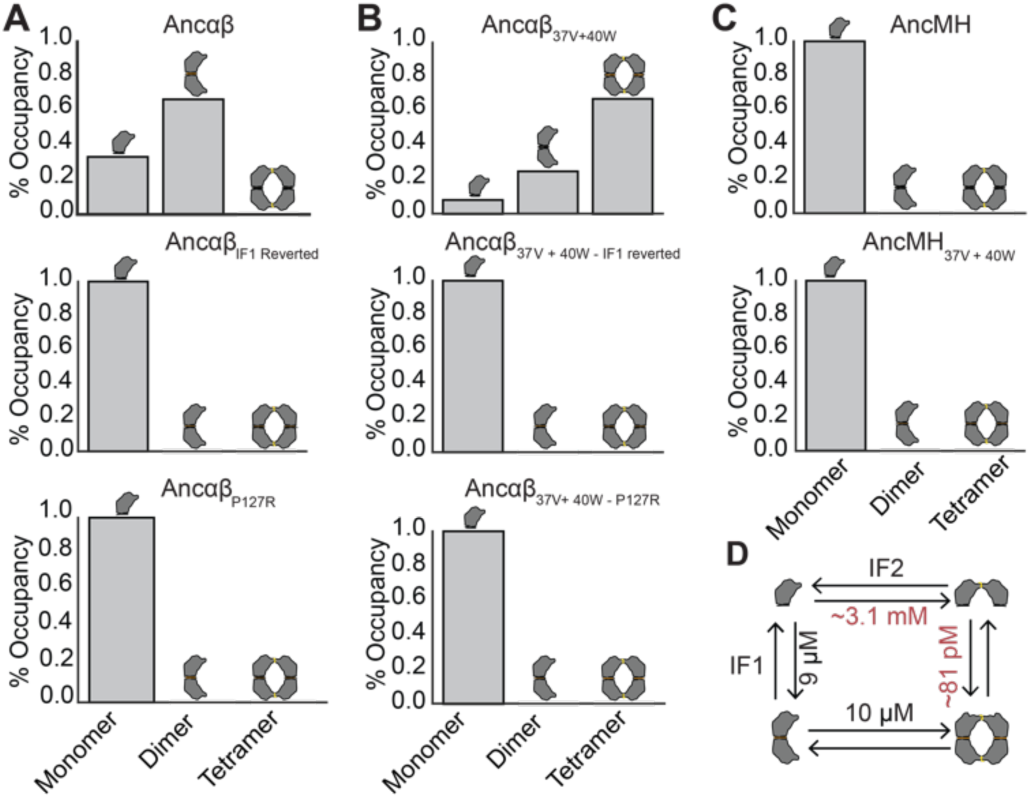
Multimerization across IF2 requires IF1. (A) IF1-mediated dimerization can be compromised by mutations. Relative occupancy of each stoichiometry as measured by native MS at at 20 mM total protein is shown for the ancestral dimer Ancɑβ (top), Ancɑβ_IF1 reverted_ (middle, a variant of Ancɑβ in which all IF1 residues are reverted to the ancestral state found in AncMH), and Ancɑβ-P127R (bottom, in which a mutation known to compromise IF1-mediated dimerization has been introduced). **(B)** Compromising IF1 prevents assembly across IF2. Relative occupancy of Ancɑβ_40W + 37V_ with and without mutations that compromise IF1-mediated dimerization. **(C)** AncMH, which does not dimerize across IF1, cannot multimerize across IF2, even when mutations sufficient to confer IF2-mediated mutimerization in Ancαβ are introduced. **(D)** Observed (black) and expected (red) affinities of Ancαβ +q40W interfaces. Expected Kd of a single iteration of IF2 (top) equals the square root of the measured apparent Kd when two iterations are present (bottom). Expected apparent Kd of two iterations of IF1 (right) equals the square of the measured Kd of a single IF1 (left).

We also tested whether assembly across IF2 could have been historically acquired before dimerization across IF1 evolved. We introduced t37V/q40W into the ancestral monomer AncMH – which existed before the evolution of dimerization -- and tested whether dimer assembly across IF2 can be conferred in this background. As predicted, only monomers were observed, with no dimers or higher stoichiometries detected (Fig. 2C). Acquisition of multimerization across IF2 by q40W and by the pair t37V/q40W therefore depends on the prior evolution of dimerization via IF1.

These observations can be explained by a simple model in which the two symmetrical interfaces contribute independently to the energy of binding. A single iteration of IF2 is too weak to confer measurable binding of two monomers into a dimer; however, IF1 is stronger and mediates assembly of dimers. Each IF1-mediated dimer presents two iterations of the IF2 surface patch, doubling the total energy of IF2-mediated assembly of dimers into tetramers. Because of the exponential relationship between energy and occupancy, a weak IF2 can therefore confer high-affinity binding but only if IF1 is already present. The affinities that we measured are consistent with this simple model. If the energy of dimer-to-tetramer assembly is twice that of monomer-dimer binding using the same interface, then the Kd of IF2-mediated tetramerization should be the square of the Kd of IF2-mediated dimerization (Fig. 2D). The Kd of the dimer-tetramer transition by Ancαβ _t37V/q40W_ across IF2 is 1 mM, which predicts that the affinity of IF2-mediated monomer-dimer transition when IF1 is compromised should be ∼ 1mM. Consistent with this prediction, we detected no dimer occupancy by Ancαβ _t37V/q40W; IF1reverted_ using an assay that can quantify Kd up to 400 µM (see Methods). This simple additive model therefore explains most – and possibly all -- of the difference in affinity conferred when IF2 is doubled in the symmetrical tetramer. The dependence of assembly across IF2 upon the presence of IF1 does not imply any direct physical interaction between the interfaces or any conformational change in one interface caused by binding at the other. We cannot rule out the possibility that IF1 binding may also allosterically modify IF2 and increase its affinity beyond the additive effect conferred by isologous repetition alone; however, any such effect must be relatively small,.

Taken together, these data indicate that the isologous architecture of IF1 and IF2 facilitated the evolution of the Hb tetramer via substitution q40W. Without this doubly symmetrical architecture, IF2 would have been too weak to mediate multimerization. The dependence of q40W’s effect on the presence of IF1 also creates contingency and order-dependence in the evolution of the Hb complex. We previously showed that IF1 evolved before the duplication of the dimeric ancestor Ancαβ (17). Our present results show that if that IF1-mediated dimer had never evolved, substitution q40W at IF2 would not have been sufficient to drive the acquisition of the tetrameric stoichiometry, and the ancestral Hb protein would have remained a monomer. If events had occurred in the opposite order – with the affinity-enhancing substitution at IF2 occurring first – this intermediate ancestor would have been a monomer; when the substitutions that confer binding across IF1 did occur, they would have triggered an immediate evolutionary transition from monomer to tetramer.

### Heteromeric specificity evolved at a single interface

We next focused on understanding the evolution of Hb’s specificity for the heterotetrameric form, which was acquired during the same phylogenetic interval after the duplication of Ancαβ. Our first question was whether specificity for heteromeric interactions was conferred by sequence changes at IF1, IF2, or both. Our previously published experiments suggest that evolutionary changes at IF2 confer no specificity: when all historical substitutions that occurred at the IF2 surface during the post-duplication interval are introduced into Ancαβ and this protein is coexpressed with Ancα, an indiscriminate mixture of homotetramers, α_1_β_3_ heterotetramers, and α_2_β_2_ heterotetramers is produced (17). We therefore hypothesized that heterospecificity of the Hb tetramer is encoded entirely by IF1, such that Ancα and Ancβ specifically heterodimerize across IF1, and these heterodimers then bind to each other via a nonspecific IF2, yielding α_2_β_2_ heterotetramers.

This hypothesis makes two predictions: 1) IF1 mediates specific assembly of α and β subunits into heterodimers, and 2) this specificity is sufficient to account for the heterospecificity of α_2_β_2_ heterotetramer. To test the first hypothesis, we characterized the specificity of hetero-vs homodimer assembly by IF1 under two different conditions in which no binding across IF2 occurs. First, we diluted a coexpressed mixture of Ancα and Ancβ to concentrations at which dimers rather than tetramers assemble: at 50 µM, only heterodimers and heterotetramers form; at 5 µM, only heterodimers are observed (Fig. 3A). IF2 does not mediate assembly of monomers into dimers in the absence of IF1 (Fig. 2A, 2B), so these heterodimers must be IF1-mediated, indicating that IF1 is heterospecific (Fig. 3A). Second, we expressed Ancα and Ancβ separately and mixed them at equal and moderate concentration; because tetramerization requires co-folding, only IF1 dimers form (33), and these are predominantly heterodimers (Fig. 3B, Fig. S5). Finally, we engineered protein Ancβ’ – a variant of Ancβ in which all IF2 residues that were substituted between Ancαβ and Ancβ are reverted to the ancestral state, thus abolishing binding across IF2– and found that it also forms predominantly heterodimers when mixed with Ancα (Fig. 3C, Fig. S5). Together, these data indicate that the derived IF1 is specific, preferentially mediating assembly into heterodimers.

**Figure 3.**
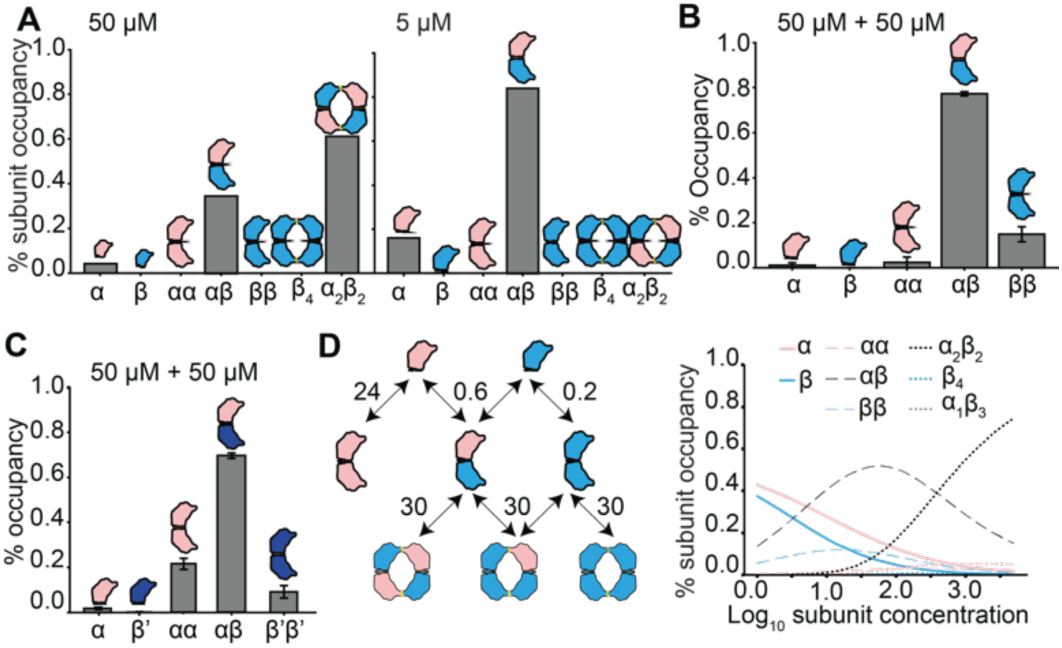
Heterotetramer specificity is conferred by specificity at IF1. **(A)** Occupancy (as fraction of all Hb subunits) when Ancɑ +Ancβ are coexpressed, measured by native MS. At 50 uM total protein, heterotetramers and heterodimers predominate (left). At 5 uM (right) – at which assembly occurs only across the high-affinity interface (IF1) -- all dimers are heterodimers. (**B)** Occupancy of subunits in stoichiometries as measured by nMS when Ancɑ and Ancβ are separately expressed and then mixed at 50 mM each; IF2-mediated tetramer assembly does not occur under these conditions, and dimers are predominantly heterodimers. Error bars represent SEM for three replicates. **(C)** Percent occupancy of stoichiometries when Ancɑ and Ancβ’ (Ancβ with all derived IF2 surface residues reverted to the state in Ancαβ) are expressed separately and then mixed at 50 uM. **(D)** Predicted occupancy of multimeric stoichiometries if IF1 is specific and IF2 is nonspecific. Left: binding scheme with experimentally estimated Kds (in mM) for IF1 and IF2-mediated multimerization by Ancα + Ancβ, assuming that all IF2 Kds are equal (for Kds, see Fig. 4D and 1D). Right: expected occupancies of each monomer, dimer, and tetramer, given the binding scheme at left. Occupancies are expressed as the fraction of all subunits in each species.

To test the second prediction – that heterospecificity mediated by IF1 is sufficient to drive specific assembly of α_2_β_2_ heterotetramers even if IF2 is nonspecific – we measured the affinities of homomerization and heteromerization across IF1 and used these measurements to predict their effects on tetramer specificity in the absence of any specificity at IF2. Using nMS and Ancβ’, we found that IF1’s heterodimerization affinity (Kd=0.6 µM) is slightly worse than its homodimerization affinity (0.2 µM), but both are far better than the Ancα homodimer (24 µM) (Fig. 3D, S6, S7, S8, S9). We then predicted occupancy of each stoichiometry as the concentration of Hb subunits changes, given these affinities at IF1 and assuming that IF2 has a dimer-to-tetramer affinity of 30 µM, as measured in Ancα + Ancβ, with no preference for homomeric or heteromeric binding (Fig. 1D). At low concentrations, the system produces almost exclusively IF1-mediated heterodimers. The predominance of heterodimers is attributable to Ancα’s weak homodimerization affinity; the excess of unbound Ancα subunits causes Ancβ subunits to preferentially heterodimerize rather than homodimerize at equilibrium, even though Ancβ’s homodimerization affinity is slightly stronger than its heterodimerization affinity (Fig. 3D). As protein concentration increases, these dimers begin to assemble with each other across IF2 into tetramers. The excess of heterodimers over homodimers means that the vast majority of the tetramers are heterotetramers, even though IF2 itself does not distinguish between subunit types. At physiologically relevant concentrations of 3mM total Hb subunits (34), the population is dominated by α_2_β_2_ heterotetramers, with a small fraction of heterodimers and virtually no homotetramers (Fig. 3d; right panel).

Taken together, these data establish that the measured specificity of IF1 alone mediates highly specific assembly of Ancα+ Ancβ into heterotetramers, even when IF2 is entirely nonspecific -- which our previous experiments suggest is the case – because IF1 is a much stronger interface than IF2. The historical acquisition of heterospecificity across IF1 after the Ancαβ gene duplication is therefore sufficient to account for the evolution of Hb’s heterotetrameric architecture.

### Heteromeric specificity evolved primarily by reducing homodimerization affinity of Ancα

Given our finding that heterospecificity evolved at the IF1 interface, we next sought to characterize whether the acquisition of specificity was driven by evolutionary changes in the α subunit, the β subunit, or both.

The heterospecificity of a pair of dimerizing proteins can be quantified in energetic terms as the difference in the ΔG of binding between the heterodimer and the mean of the two homodimers (ΔΔG_spec_; see methods for calculation). If ΔΔG_spec_ = 0, then the fractional occupancy of the heterodimer at saturating and equal concentrations of subunits will be 50%, as will the sum of the homodimers; this is true even if the homodimer ΔGs are very different from each other, as long as the heterodimer ΔG is halfway between them. By contrast, if ΔΔG_spec_<0, then heterodimers will account for the majority of dimers; conversely, if ΔΔG_spec_>0, homodimers together will predominate (Fig. 4A-C). Hetero- or homospecificity thus arises when two paralogs contribute nonadditively to dimerization. Whether or not the system is hetero- or homospecific, the two homodimers will have equal occupancies to each other at saturating conditions: irrespective of the fraction of subunits assembled into heterodimers, the remaining subunits will be at equal concentrations and by definition will be well above the homodimerization Kds (28).

**Figure 4.**
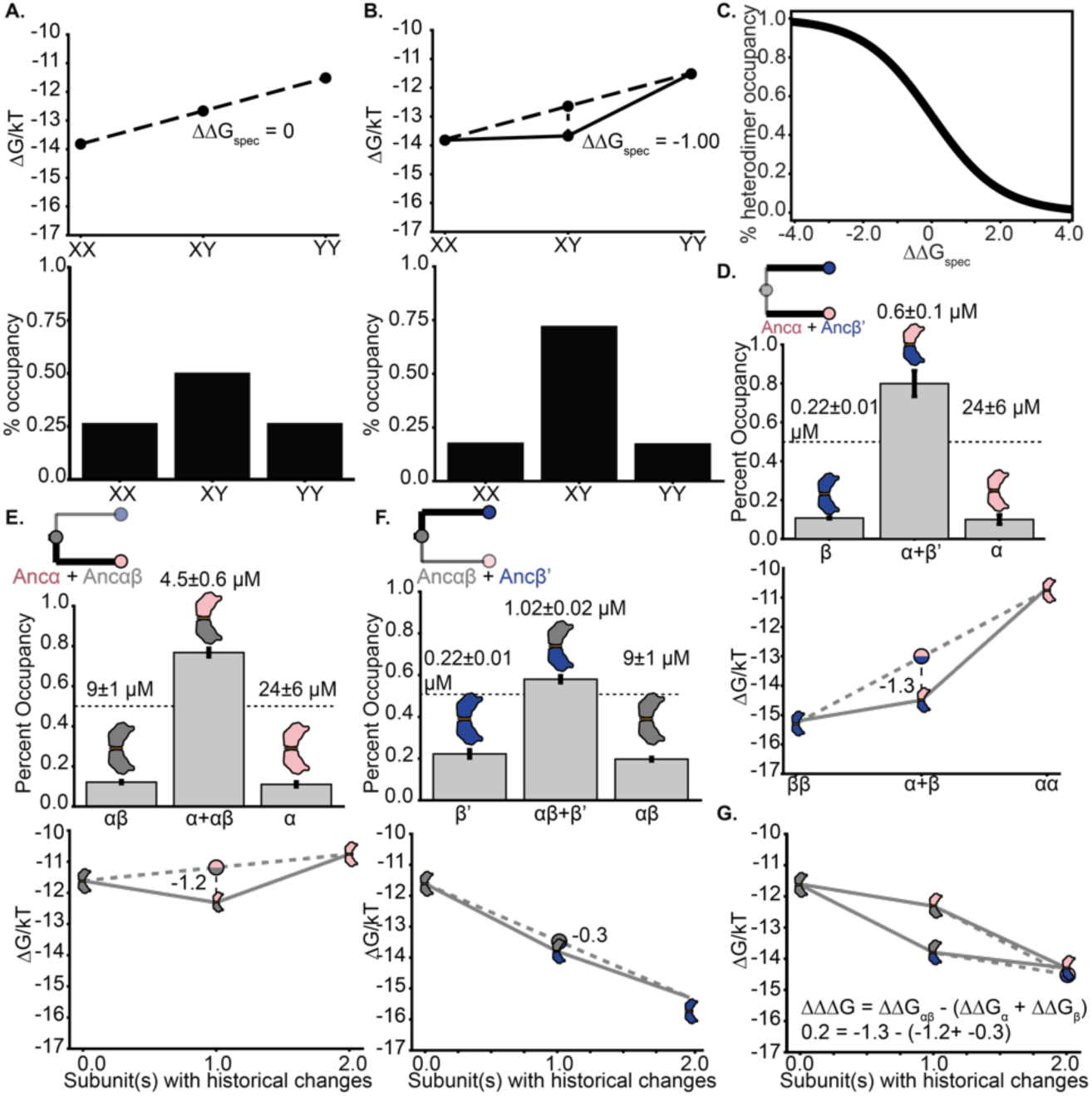
Contribution of historical changes in each subunit to the acquisition of heterospecificity. **(A)** Theoretical example of affinities and occupancy in a system of dimers with no specificity. *Top*: ΔG of dimerization for homodimers (XX and YY) and heterodimers (XY), in units of kT. In the absence of specificity, ΔG of the heterodimer equals the average of the homodimers (dotted line). *Bottom*: expected fractional occupancies of dimers at 1 mM per subunit and dissociation constants (Kd), given the ΔGs in the top panel. In the absence of specificity, heterodimer occupancy = 50%. **(B)** Example of a system with preference for the heterodimer. ΔΔG (the deviation of the heterodimer ΔG from the average of the homodimers) is shown. *Bottom*: Kd and predicted occupancy of each dimer at 1 mM. **(C)** Relationship between ΔΔG and heteromeric occupancy at 1 mM per subunit, assuming the ΔGs of homodimerization for as shown in panel A. **(D)** Specificity of IF1 dimerization in system of Ancα+Ancβ’. *Top*: expected fractional occupancies at 1 mM, given Kds assessed by nMS (shown above each bar, with 95% confidence interval). *Bottom*: ΔGs and ΔΔG given measured Kds. Dotted line, expected occupancies in the absence of specificity. **(E)** Specificity of IF1 acquired on the branch leading from Ancɑβ to Ancα, shown as occupancy and ΔGs of the Ancɑβ + Ancɑ system. The number of subunits that contain historical changes in each dimer is shown relative to the Ancɑβ homodimer. **(F)** Specificity of IF1 acquired on the branch leading from Ancɑβ to Ancβ, shown as occupancy and ΔGs of Ancɑβ + Ancβ’. **(G)** Interaction effect on specificity when evolutionary changes leading from Ancαβ to Ancα (pink) and Ancβ’ (blue) are combined. Homodimer of Ancαβ (gray) and each heterodimer are plotted by their ΔG. The observed ΔΔG of each heterodimer in combination Ancαβ is shown (see panels D-F). If the specificity acquired in the two subunits affects heterodimerization independently, then ΔΔG of Ancα+Ancβ will equal the sum of the ΔΔGs, yielding a parallelogram. The deviation from this expectation is shown.

We quantified the heterospecificity of Ancα and Ancβ at IF1 by estimating ΔΔG_spec_. We used nMS to measure the homodimer and heterodimer affinities of Ancα and Ancβ’, which contains all substitutions that occurred along the Ancβ branch except those that mediate tetramerization across IF2, allowing us to prevent tetramerization and thus isolate the specificity effects at IF1. From these affinities, we calculated the ΔG of binding and the expected fractional occupancy of each dimer at high and equal concentration of subunits. For Ancα+Ancβ’, we found that ΔΔG_spec_= –1.3 (in units of kT) and heterodimer occupancy of 82% (Fig. 4D). This represents the total specificity acquired by the two diverging paralogs after the duplication of of Ancαβ, which by definition had no specificity. This specificity was acquired because of evolutionary changes in all three relevant affinities. Relative to the ancestral dimerization affinity of Ancαβ, Ancα‘s energy of homodimerization became worse (ΔΔG = 0.9) while homodimerization by Ancβ’ improved substantially (ΔΔG = –3.7). The heterodimer affinity improved by ΔΔG _=_ –2.7, substantially more than the average of the two homodimers, yielding the observed strong preference for the heterodimer.

We next sought to isolate the contribution of the evolutionary changes that occurred along each of the two branches. To measure the specificity acquired along the branch leading to Ancα, we measured affinities and calculated ΔΔG_spec_ when Ancα is mixed with the deeper ancestor Ancαβ. This pair of proteins is heterospecific, with ΔΔG_spec_= –1.2 (expected heterodimer occupancy 76%). Changes in the α subunit alone therefore account for >90% of the total ΔΔG_spec_ that was acquired by the entire Ancα+Ancβ’ system. This specificity is acquired via a 2.6-fold reduction in Ancα’s homodimerization affinity compared to the Ancαβ ancestor and a 1.8-fold improvement in heterodimer affinity (Fig. 4E; Fig. S7B & D).

To isolate the contribution to IF1 specificity of evolutionary changes that occurred along the branch to Ancβ, we measured affinities when Ancβ’ is mixed with Ancαβ. This pair of proteins is weakly heterospecific, with ΔΔG_spec_*=* –0.3 and expected heterodimer occupancy of just 58%. The specificity is weak because both the homodimer and heterodimer affinity improved, but the deviation of the heterodimer from the average of the homodimers is very small (Fig. 4F; Fig. S7A&C).

Finally, we assessed whether the evolutionary changes in the Hbα subunit and those in the Hbβ subunit interacted with each other nonindependently. If the changes affect specificity entirely independently, ΔΔG_spec_ should equal the sum of the ΔΔG_spec_ acquired on each of the two branches (–1.2 + –0.3 = –1.5). The observed ΔΔG_spec_ = –1.3, suggesting a very weak negative interaction between changes in the two subunits, which makes the complex slightly less heterospecific than expected if the substitutions were independent (Fig. 4G).

Taken together, these data indicate that the IF1 specificity acquired by the derived complex Ancα + Ancβ is primarily attributable to substitutions in the α subunit, with substitutions in the β subunit making a much smaller contribution; nonadditive interactions between the two sets of changes did not contribute to the evolution of heterospecificity. The most important factor was that Ancα became much worse at binding itself than at binding Ancβ. Ancβ, by contrast, became slightly worse at binding Ancα than binding itself (Fig. 4G).

### A one-residue deletion was the primary evolutionary cause of heterospecificity

We next sought to identify the particular historical substitutions in Ancα that conferred this heteromeric specificity on IF1. Only three sequence changes occurred on the branch from Ancαβ to Ancα: a single-residue deletion of a histidine at site 3 (ΔH3), a five-residue deletion in helix D (ΔD), and an amino acid replacement (v140A). ΔH3 is on the protein’s N-terminal loop near IF1, and ΔD directly contributes to the interface. Substitution v140A is biochemically conservative and far away from the interface. The deletions are conserved to the present in Hbα subunits throughout the jawed vertebrates, including humans, whereas the amino acid at site 140 varies (Fig. S1). We therefore focused first on the effects of the deletions.

To isolate the contribution of each deletion to the evolution of specificity, we introduced each one singly into Ancαβ and measured its effect on affinity and specificity when the mutant protein is mixed with Ancαβ. We found that introducing ΔH3 alone confers substantial specificity, recapitulating >80% of the Ancα’s acquired heterospecificity for Ancαβ (ΔΔG_spec_ = –1.0 out of a total ΔΔG_spec_ = –1.2 acquired along this branch) and >75% of the specificity acquired along both branches by the entire Ancα+Ancβ’ complex (ΔΔG_spec_ = –1.3, Figs. 5A, C). ΔH3 enhances specificity by improving heterodimer affinity and reducing homodimer affinity, with both Kds very close to those of Ancα (Fig. 5A; Fig. S10A & C).

**Figure 5.**
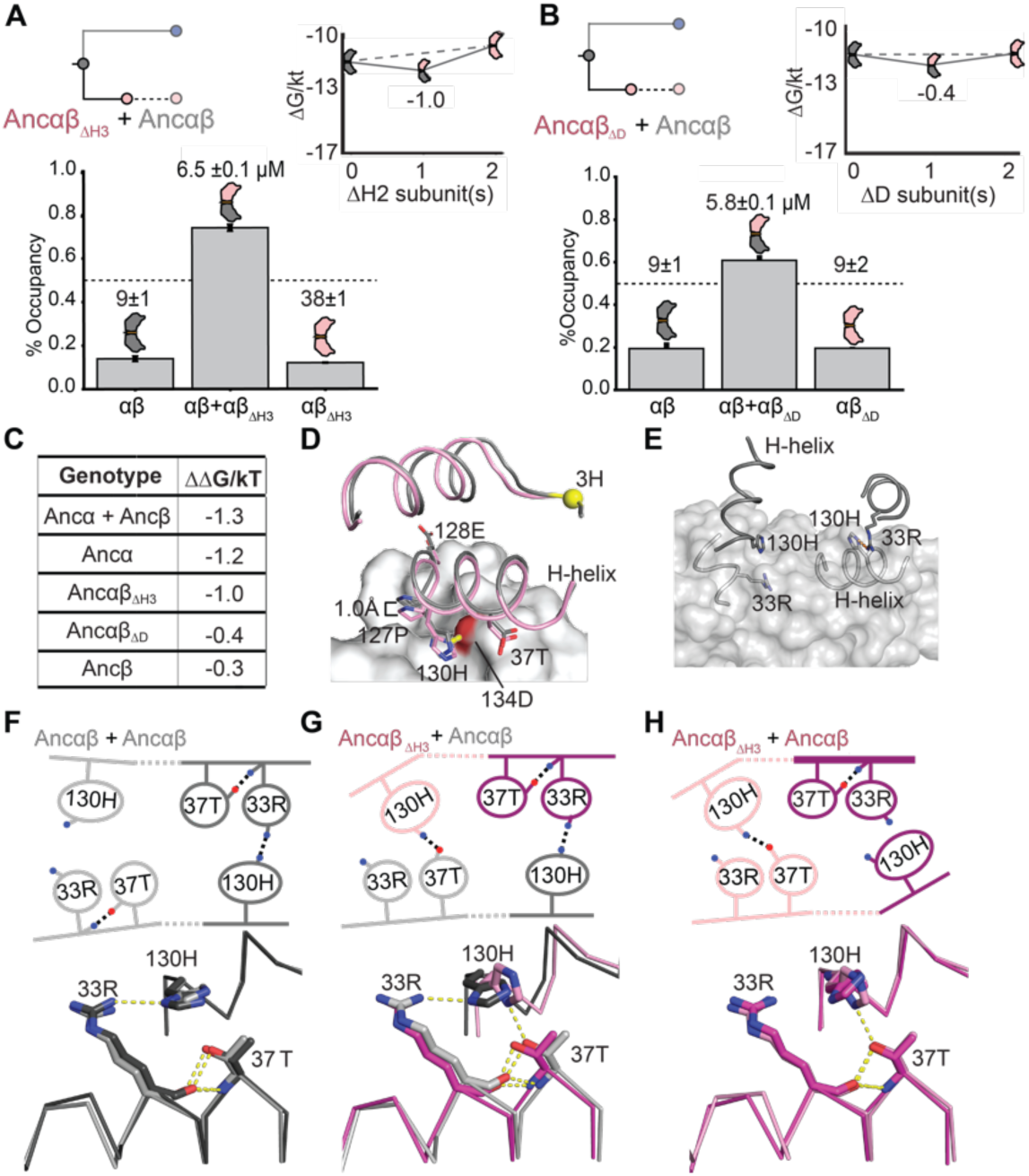
Effect of historical sequence changes on specificity. **(A)** Specificity of Ancɑβ_ΔH3_ with Ancɑβ, as in Fig. 4. **(B)** Specificity of Ancɑβ_ΔD_ with Ancɑβ. **(C)** Gain in specificity caused by various sets of historical mutations, relative to Ancɑβ. Ancɑ+Ancβ, all changes on both post-duplication branches. Ancɑ, all changes on the branch leading to Ancɑ. ΔΗ2 and ΔD, deletions that occurred on the Ancɑ branches. **(D)** Models of Ancɑβ homodimer and Ancɑβ_ΔH3_ + Ancɑβ heterodimer. The N-terminal helix and the portion of IF1 involving helix H is shown. Grey surface, Ancɑβ subunit common to both models. Grey cartoon, other Ancɑβ subunit in the homodimer; pink cartoon, Ancɑβ_ΔH3_ subunit in the heterodimer. Yellow, 3H residue deleted in Ancɑβ_ΔH3_. Helix H side chains in the interface are shown as sticks. The hydrogen bond in the heterodimer from 130H to 37T (red surface) is shown (dotted line). **(E)** A portion of IF1 in the Ancɑβ homodimer model, showing the isologous interactions with imperfect symmetry between 130H and 33R. Orange dashed-line, hydrogen bond. The two subunits are colored different shades of gray. The surface of the light-gray subunit is shown. **(F, G, H)** Key residues in IF1 with hydrogen bonds that are affected by ΔΗ3 in the homodimers and heterodimer of Ancɑβ and Ancɑβ_ΔH3._ Top, cartoon of key contacts. The two iterations of these interactions across the isologous interface are shown, one each in light or dark hue. Blue and red, nitrogen and oxygen atoms, respectively. Dotted lines, hydrogen bonds. The change in position of the H-helix caused by ΔΗ2 is shown. Bottom, structural alignment of the two iterations of the isologous interface in each dimer. Each dimer structure was duplicated exactly and then aligned to the original by targeting one subunit of the copy to align to the other subunit of the original. Hues correspond to the isologous iterations in the cartoon above.

The other deletion, ΔD, removes several residues that directly interact with the other subunit across IF1, but introducing this change into Ancαβ had a much weaker effect on specificity (ΔΔG_spec_ = –0.4, Fig. 5B; Fig. S10B & D). When the contributions of ΔH3 and ΔD to specificity are added together, they slightly exceed the specificity of Ancα (–1.4 rather than –1.3), suggesting the possibility of a very weak negative epistatic interaction between them or a small countervailing effect of the third change v104A. Taken together, these results indicate that ΔH3 was a large-effect historical sequence change that accounted for most of the specificity historically acquired by the derived Hb complex.

### Structural mechanisms for the gain in specificity

We next considered the structural mechanisms by which ΔΗ3 conferred specificity by increasing heterodimer affinity and reducing homodimer affinity. For a mutation to have these opposite effects, it must yield favorable interactions when introduced into one side of the interface (in the heterodimer) but have deleterious effects when introduced twice (in the homodimer). In principle, two kinds of mechanisms could cause these opposite effects. Either 1) the mutated residue could interact directly with the same residue on the other subunit favorably when one is in the derived state but unfavorably when both are, or 2) the symmetry of the interface could be imperfect, such that introducing the mutation on one side of the interface is favorable but introducing it again onto the other side is net-unfavorable. The first scenario does not pertain in this case. Residue H3 is part of the N-terminal loop, which does not participate directly in IF1 but instead packs against helix H, which does contribute to IF1. But neither helix H nor the N-terminal loop contact the same elements in the other subunit across the interface (Fig. 5D). Asymmetry in the interface is therefore the likely of cause ΔΗ3’s differential effects on heterodimer vs. homodimer affinity.

To gain insight into the possible nature of this asymmetry and potential mechanisms by which ΔH3 affects specificity, we modeled the structures of the Ancαβ homodimer, the Ancαβ_ΔΗ2_ homodimer, and the heterodimer of these two proteins. The modeled Ancαβ homodimer contains a subtle asymmetry: on one end of IF1, residue 130H on helix H sits close to 33R on the opposite subunit, which allows a cross-interface hydrogen bond to form; on the other end of the interface, the two residues are slightly further away from each other, leaving their hydrogen-bonding potential unsatisfied when bound (Fig. 5E & F). In the heterodimer, deleting Ηis2 from one subunit repairs this unfavorable interaction. Specifically, the deletion shortens the N-terminal loop and changes its packing interaction against helix H, which causes helix H to slide along the interface by ∼1 Å compared to its position in the unmutated Ancαβ homodimer (Figs. 5D, 5G). 130H moves closer to 37T on the other subunit, allowing it to form a new hydrogen bond across the interface, and several other interactions across the interface are also enhanced. On the other end of the isologous interactions, the favorable interactions found in the homodimer remain intact. This provides a potential structural explanation for how ΔΗ3 improves heterodimer affinity (Figs. 5D, G).

The modeled Ancαβ_ΔΗ3_ homodimer structure is notably asymmetric and suggests a possible mechanism by which introducing ΔΗ3 into both subunits reduces affinity (Fig. 5H). One side displays the favorable new cross-interface interactions caused by ΔΗ3 in the heterodimer, including the 130H-37T hydrogen bond. On the other side, however, the effect of the deletion is very different: ΔH3 again causes helix H to slide along the interface, but on this side the movement of 130H breaks the ancestral 130H-33R hydrogen bond, and 37T is also too far away to interact favorably. This leaves the side chains of both 130H and 33R unsatisfied, reducing homodimer affinity. In total, the homodimer of Ancαβ_ΔΗ3_ contains three unsatisfied hydrogen-bond donors/acceptors at these sites, whereas only one and two are unsatisfied in the heterodimer and the ancestral homodimer, respectively.

These hypothesized mechanisms appear to have persisted over time. The same pattern of interactions are found in the modeled structures of the hetero- and homodimers of Ancα + Ancβ (Fig. S11). High-resolution crystal structures of extant hemoglobin also show notable asymmetries in the multimerization interfaces, which exceed the deviation expected given the resolution of the structures (35). These structures include some of the particular asymmetrical interactions observed in our ancestral models: in the human Hb heterotetramer, 33R hydrogen bonds across IF1 to residue 130, but this interaction is again lacking in the homodimer of human Hbα, leaving 33R unsatisfied, potentially explaining the weak homomeric affinity of Hbα (Fig. S11). At least some of the mechanisms of heterodimer specificity suggested by the structural models of the ancestral proteins are therefore present in the known structures of its present-day descendants. Structural models are prone to error, and the asymmetries we observed are subtle; further research will be required to definitively characterize potential asymmetries in the ancestral multimers.

### Multiple historical sets of substitutions could have conferred heterospecificity

If specificity in an isologous interface can evolve simply by causing nonadditive impacts on the binding energies of heterodimer and homodimers, then there should be many mutations that have the potential to make the interface specific in one direction or another. Indeed, if the interface’s symmetry is imperfect, then most mutations that affect affinity should impart specificity to some degree.

To test this hypothesis, we measured the effect on specificity of subsets of changes that occurred along the Ancβ lineage, which the results above show had strong effects on affinity when introduced all together. First, we tested the five substitutions that that occurred at the IF1 surface (Fig. 5E & 5F). We introduced these changes into Ancαβ (creating protein Ancαβ_IF1_) and measured affinity and specificity when this protein is mixed with Ancαβ. These substitutions yield a highly heterospecific complex (ΔΔG_spec_ = –2.2, heterodimer occupancy 90%, Fig. 6A; Fig. S12A, C, & E). Unlike the Ancα substitutions, the Ancβ_IF1_ substitutions confer heterospecificity by improving both homodimer and heterodimer affinity, but they improve the latter by more than the former.

**Figure 6.**
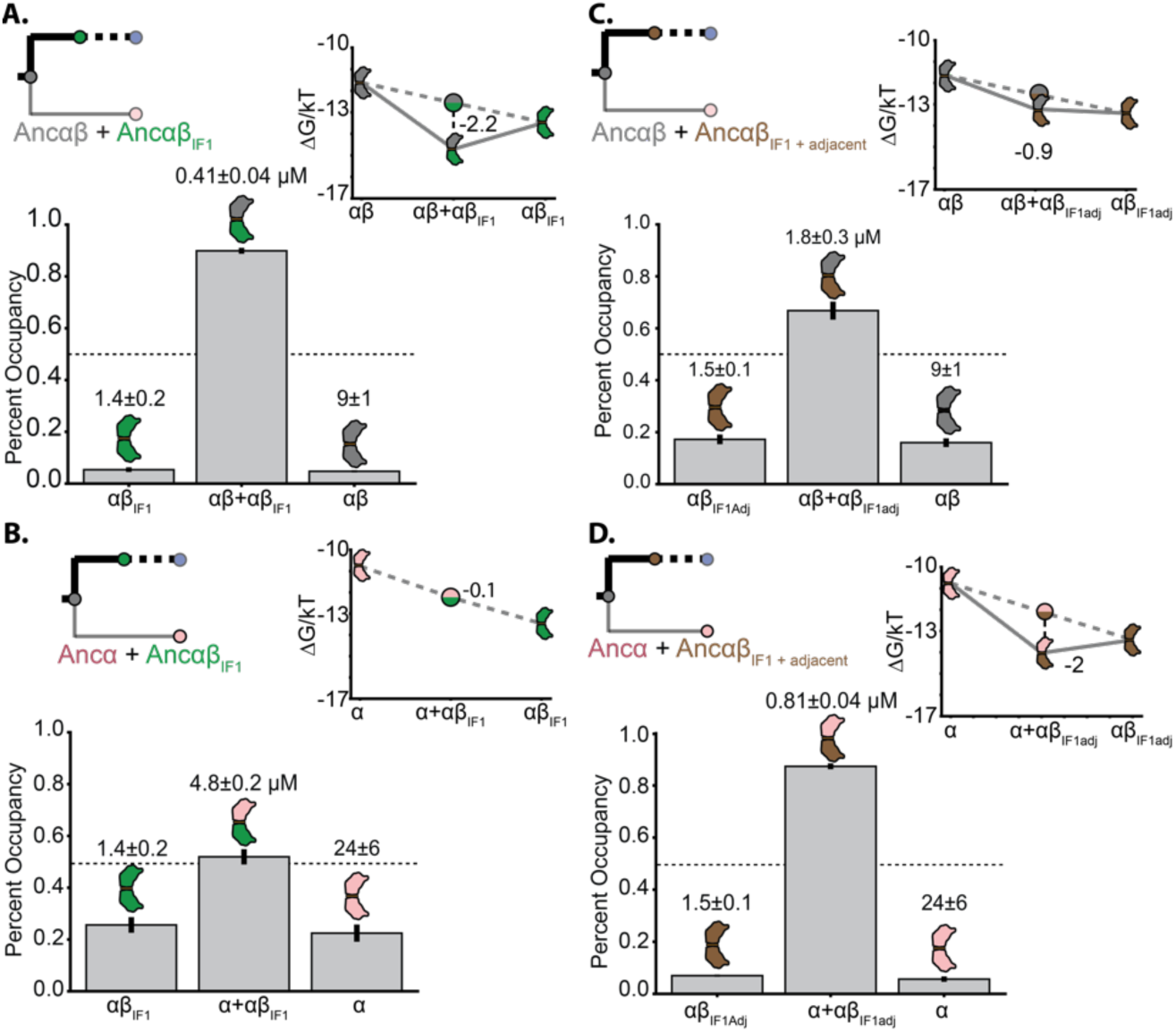
Other subsets of historical substitutions confer heterospecificity on IF1. Affinities measured by nMS, predicted occupancy based on those Kds at 1 mM each subunit, and ΔΔG_spec_ are shown for A) Ancɑβ + Ancɑβ_IF1_, which contains the five substitutions at the IF1 surface that occurred in the Ancβ lineage; B) Ancɑβ + Ancɑβ_IF1 + adjacent_, which also includes 4 additional substitutions in Ancβ near but not on the interface; C) Ancɑ + Ancɑβ_IF1_, and D) Ancɑ + Ancɑβ_IF1 + adjacent_.

Because Ancαβ_IF1_ is specific in complex with Ancαβ, we wondered whether it would also be specific with Ancα. We found that this complex is barely heterospecific (ΔΔG_spec_ = –0.1, Fig. 6B), implying that other substitutions on the branch leading to Ancβ but not on the interface must have contributed to the evolution of specificity between Ancαβ_IF1_ and Ancα. We therefore introduced an additional set of five historical substitutions that occurred in Ancβ but one structural layer away from IF1 (see ref. 17). This protein (Ancαβ_IF1+Adjacent_) has strong heterospecificity when mixed with Ancα (ΔΔG_spec_ = –2.0, heterodimer occupancy >85%, Fig. 6D; Fig. S12B, D, & F), because these mutations together improve both heterodimer and homodimer affinity, but with a larger improvement in the heterodimer. It is also moderately heterospecific when mixed with Ancαβ (ΔΔG_spec_ = –0.9).

Finally, we tested the effect of the adjacent substitutions on their own and found that they confer specificity when mixed with Ancαβ (ΔΔG_spec_ = –0.9). These mutations impart specificity by causing almost identical changes in homo- and heterodimer affinity. They also confer some heterospecificity when Ancαβ_Adjacent_ is mixed with Ancα (ΔΔG_spec_ = –0.6, Fig. S13A-D).

There are therefore several distinct sets of substitutions that occurred during history, and which can be sufficient to confer heterospecificity on their own (and in various combinations), and they do so via distinct patterns of effects on affinity. This degeneracy of mechanisms for evolving specificity arises because there are many ways in which the energy of binding can change nonadditively between heterodimer and homodimer. In every case, heteromeric specificity rather than preference for the homomer was the result.

## DISCUSSION

This work provides a mechanistic history of the evolutionary transition from the ancestral Ancαβ homodimer to the derived Hb heterotetramer (summarized in Fig. S1). Each transition was driven by a very simple genetic mechanism: a single substitution at IF2 conferred high affinity tetramerization, and a single amino acid deletion at IF1 conferred heteromeric specificity. These two key sequence changes have remained conserved in the descendant Hbα and Hbβ subunits of all extant jawed vertebrates (Fig. S1). These transitions were both facilitated by the isologous architecture of Hb’s two interfaces, which creates a propensity for mutation to produce high-order complexes and heterospecificity.

### Symmetry facilitated evolution of the tetrameric stoichiometry

We found that tetramerization across IF2 was driven primarily by a single replacement to a bulky hydrophobic amino acid (q40W). In biochemical studies of extant protein interfaces, much of the free energy change in protein-protein binding is attributable to interactions of bulky hydrophobic residues with hydrophobic surface indentations (36), and mutations to bulky hydrophobic amino acids can drive assembly into high-order multimers (9,37–40). Similar substitutions during history may have been driving mechanisms during the evolution not only of Hb but of other molecular complexes, as well.

The majority of complexes assemble through isologous interfaces (41). It has been suggested that this must imply that isology confers some selective benefit by improving protein function (1). Our results suggest an alternative explanation. If mutations are much more likely to produce isologous complexes than nonisologous ones, then isologous complexes will predominate in nature, even if there is no systematic fitness difference between the two types of multimer. We found that although IF2 is intrinsically weak and mutation q40W cannot confer dimerization on its own, it can drive tetramerization if its effects are multiplied in an isologous higher-order complex. By contrast, If the interfaces were non-isologous -- with q40W interacting with a hydrophobic divot on some other surface of the facing subunit – then this favorable interaction would appear only once, and it would be insufficient to substantially improve binding energy and confer meaningful tetramer occupancy. By this explanation, isologous complexes are abundant because they are easier to produce by mutation than head-to-tail multimers, not more likely to be fixed by selection.

It has been observed that in high-order multimers, the interface with higher affinity usually evolves before the lower-affinity interface(s) (42–45). Hb evolution displays this pattern, with the stronger interface IF1 evolving before IF2 (17). It has been suggested that this pattern is attributable to selection: by this hypothesis, selection favors evolutionary intermediates that contain the high-affinity interface, because those containing only the low-affinity interface assemble slowly and/or misassemble into anomalous complexes (42,43). Our work here suggests a different explanation: it is easier for mutations to generate a new interface that confers a higher stoichiometry if a strong interface is already present, because the affinity of the new interface is multiplied by iteration in an isologous complex. By contrast, low-affinity interfaces do not confer multimerization on their own, so if the low-affinity interface were to evolve first, then the effects of mutations on the second interface would not be multiplied. In complexes with multiple interfaces, the stronger interface tends to be older not because such trajectories improve fitness but because mutation is more likely to build elaborate complexes in this historical order.

### One interface confers specificity on a higher-order multimer

Our experiments show that evolutionary change at just one of Hb’s interfaces was sufficient to confer specific assembly into heterotetramers. Specificity at IF1 alone was sufficient to mediate the heterospecificity of the tetramer because this interface is so much stronger than IF2: IF1 mediates the specific assembly of heterodimers, which assemble into heterotetramers across IF2, even though IF2 itself confers little or no specificity.

The specificity of IF1 and the isology of the complex also explains the *trans* conformation of Hb’s quaternary structure, in which each Hbα subunit binds one Hbβ subunit across IF1 and a different Hbβ across IF2. The alternative *cis* conformation -- in which Hbα is paired with an Hbα (and Hbβ with Hbβ) across one of the interfaces – is never observed. Although IF2 imposes little or no specificity, its isologous orientation necessarily means that the two IF1-mediated heterodimers must be rotated 180° relative to each other, placing each Hbα across IF2 from the Hbβ of the other heterodimer. In the *cis* conformation, the heterodimers would not be rotated 180° relative to each other, and all the favorable interactions that IF2 comprises would not form; residue 40W, for example, would not face the hydrophobic divot on IF2 across the interface. Given the heterospecificity of IF1, isology constrains the Hb tetramer to its *trans* α_2_β_2_ architecture.

These observations suggest a simple and potentially general mechanism for the evolution of specificity in the quaternary structures of high-order multimers. Specificity need not evolve at every interface in the complex, especially if the interfaces are isologous. Rather, mutations need only make the stronger interface specific to confer assembly into particular high-order architectures.

### Symmetry allowed specificity to evolve in one subunit

We found that a single genetic change in one paralog – a one residue deletion in Ancα -- was sufficient to confer IF1’s heterospecificity. This result contrasts with prior studies of nonisologous complexes, in which heterospecificity evolved because of genetic changes in both interacting subunits (7,13,18,19,21,24–26).

This difference in historical genetic mechanism reflects the opportunities presented by the two different types of multimeric architecture. In nonisologous complexes, a mutation in the “head” of one duplicate gene will not be sufficient to distinguish between its own tail and that of its paralog (unless it somehow changes the conformation of both distinct surfaces). In an isologous complex like Hb, however, a change in one subunit can confer specificity, because it makes the interface different between the heterodimer, the mutated homodimer, and the unmutated homodimer.

Acquiring specificity in an isologous interface requires the mutation to nonadditively change the affinity of the heterodimer relative to the homodimers. If the symmetry of such interfaces were perfect, a mutation in one subunit would affect interactions across the interface identically on each side of the interface, resulting in additive effects on affinity. Nonadditivity would arise only if mutations affect sites that interact with each other across the rotated interface. This would require either a mutation at the precise axis of rotational symmetry or multiple mutations at several sites.

If the symmetry is imperfect, however, a single mutation can affect interactions differently when it appears twice in the homodimer versus when it occurs once in the heterodimer. Imperfect asymmetry could facilitate the evolution of specificity in many complexes. Virtually all isologous interfaces contain subtle asymmetries (46). This imperfection arises for two reasons: perfect symmetry is entropically unfavorable, and amino acids near the axis seldom face each other with perfect symmetry, because each amino acid itself is asymmetrical, and this asymmetry propagates elsewhere in the interface (46,47). Extant human hemoglobin is one of many examples of isologous interfaces in which asymmetry is imperfect (35,48). Isologous interfaces therefore provide a starting point for homo- or heterospecificity to be acquired by substitutions in a single subunit.

### Specificity evolved through a single mutation

We found that a single mutation – deletion of residue His2 in the alpha subunit – conferred most of the heterospecificity of Ancα + Ancβ. This simple mechanism was possible because only a small change in relative binding energy is required to yield substantial changes in specificity. The IF1 of Ancα+Ancβ’ mediates 80% heterodimer occupancy at equal and saturating concentrations, but its specificity is quite moderate ΔΔG_spec_ = –1.3). This difference in binding energy is less than that associated with a typical hydrogen bond or burial of a large hydrophobic residue. The structural differences in physical interactions across the homodimer vs. heterodimer interfaces in our modeled structures could easily yield energetic differences of this magnitude, although the particular form of asymmetry in ancestral hemoglobin complexes remains uncertain. Recent in silico work also found that small differences in ΔG can cause large differences in occupancy between homodimers and heterodimers (28).

Why do such subtle differences in energy have such large impacts on specificity? Mutations that cause a modest deviation in binding energies can cause large changes in occupancy because of the nonlinear Boltzmann relationship between these quantities (Fig. 4C). Moreover, specificity is determined by the deviation from additivity in the heterodimer relative to the homodimers, so small differences in the free energy of binding will propagate into even larger changes in specificity. Because of this intrinsic sensitivity, we predict that the evolution of specificity in paralogous complexes with symmetrical interfaces will often be attributable to one or a few genetic changes with relatively small energetic effects and subtle structural mechanisms. That specificity can evolve so easily also implies that paralog interference after gene duplication (49,50) may often be easily resolved through one or a few mutations.

If specificity can be acquired by small deviations from energetic additivity in either direction, one might expect that homomeric and heteromeric specificity would be equally likely to evolve. But empirical observations suggest that heteromers evolve much more frequently after gene duplication (12,13). Our findings suggest a plausible explanation for this pattern. The modeled structures suggest that the critical mutation for conferring specificity on Hb does so because imperfect asymmetry in the interface creates a kind of antagonistic pleiotropy: a favorable interaction occurs when the mutation is introduced once in the heteromer, but it fails to produce the same favorable contact and even disrupts a different favorable contact when introduced again on the other side of the interface in the homomer. Heterospecificity will result whenever asymmetry causes antagonistic pleiotropy like this, such that a favorable interaction can be optimized when it is iterated once but not twice. In contrast, homomeric specificity requires a mutation to be even more favorable the second time it is introduced on the other side of an interface. For this to occur, imperfect symmetry must synergistically enhance the interactions caused by the two iterations of the mutation in the homodimer. This scenario seems far less likely than an antagonistic effect, because favorable interactions are constrained in many ways, requiring fairly precise compatibility of polarity, size, angle, etc. The imperfect symmetry of isologous interfaces may therefore create a mutational propensity that favors the evolution of heteromeric over homomeric specificity.

Taken together, our observations contribute to a growing body of evidence that complex multimeric complexes can evolve through simple genetic mechanisms (5, 15, 37, 39,51–55). In Hb evolution, a single substitution in one of the duplicated genes was sufficient to cause a doubling in stoichiometry from dimer to tetramer, and a single-residue deletion at one interface in the other subunit was sufficient to confer strong preference for the α_2_β_2_ heterotetrameric form. Although other substitutions enhanced these effects, and others may have permitted or entrenched them (5,56), our data indicate that discrete evolutionary increases in complexity can occur by very short mutational paths from simpler ancestral forms. The major effects of these small sequence changes was possible because they took place in the context of an isologous complex, and it is likely that its symmetry was slightly imperfect. Many multimers share these structural properties, so we predict that, when other multimeric complexes are studied in detail, simple mechanisms will be found to have driven their historical elaboration.

## MATERIALS AND METHODS

### Sequence data, alignment, phylogeny, and ancestral sequence reconstruction

The reconstructed ancestral sequences used here are the same as those reported previously (17). Briefly, 177 amino acid sequences of hemoglobin and related paralogs were collected and aligned. The maximum likelihood (ML) phylogenetic tree was inferred using the AIC best-fit model, LG+G+F (57,58). The phylogeny was rooted using as outgroups neuroglobin and globin X, which are found in both deuterostomes and protostomes and diverged prior to the gene duplications that produced vertebrate myoglobin and the hemoglobin subunits. Ancestral sequence reconstruction was performed using the empirical Bayes method (62), given the alignment, ML phylogeny, ML branch lengths, and ML model parameters. Reconstructed ancestors that were used in this study have been deposited previously in GenBank (IDs MT079112, MT079113, MT079114, MT079115).

The historical mutations that we introduced into those ancestral proteins are the following. For the set *IF1-reverted*, all sites in IF1 that were substituted on the branch leading to Ancb are reverted to the ancestral state found in Ancab; the mutations introduced are V36t, Y38h, V115a, V119e, H130r, D134e. For the set *IF2-reverted*, all sites that were substituted in IF2 on the branch leading to Ancb are reverted to the ancestral state found in Ancab; the mutations introduced are T37v, W40q, R43t, H100r, E104h. For the set *IF1*, all sites at IF1 that were substituted between Ancab and Ancb are changed to the derived state found in Ancb; the mutations introduced are t37V, k58M, r107K, h130Q, d134Q4. For the set *Adjacent*, five sites adjacent to IF1 that were substituted between Ancab and Ancb are changed to the derived state found in Ancb; the mutations introduced are h47S, s60N, q62K, a96S, h97E. The set *IF1+Adjacent* is the union of the sets *IF1* and *Adjacent*. Deletion DD deletes residues a54, e55, a56, i57, and k58 from Ancab.

### Recombinant protein expression

Coding sequences for reconstructed ancestral proteins were optimized for expression in *Escherichia coli* using IDT Codon Optimization and synthesized *de novo* as gBlocks (IDT). Coding sequences were cloned by Gibson assembly into vector pLIC (3) under control of a T7 polymerase promoter. For co-expression of Ancα+Ancβ, a polycistronic operon was constructed under control of a T7 promoter and separated by a spacer containing a stop codon and ribosome binding site, as described in (63).

BL21 (DE3) *Esherichia coli* cells (New England Biolabs) were heat-shock transformed and plated onto Luria broth (LB) containing 50 ug/mL carbenicillin. For the starter culture, a single colony was inoculated into 50 mL of LB with 1:1000 dilution of working-stock carbenicillin and grown overnight. 5 mL of the starter culture were inoculated into a larger 500-mL terrific broth (TB) mixture containing the appropriate antibiotic concentration. Cells were grown at 37° C and shaken at 225 rpm in an incubator until they reached an optical density at 600 nm of 0.6-0.8.

For expression of single globin proteins, 100 uM of isopropyl-β-D-1-thiogalactopyranoside (IPTG) and 25 mg/500 mL of hemin were added to each culture. Expression of single proteins in culture were done overnight at 22° C. Cells were collected by centrifugation at 4,000*g* and stored at -80° C until protein purification. Coexpressed proteins were induced using 500 mM IPTG expression with 25 mg/500 mL hemin for 4 hours at 37°C. Cells were collected by centrifugation at 4,000*g*, immediately followed by purification.

Human hemoglobin was bought commercially (Sigma-Aldridge) and resuspended in PBS.

We attempted to co-express and purify Ancαβ_ΔH3_ in complex with Ancαβ_40W_, but we were not able to identify conditions at which the two species could be expressed and purified to near-equal concentrations.

### Protein purification by ion exchange

All singly expressed proteins (all ancestral globins except Ancα+Ancβ) were purified using ion exchange chromatography. All buffers were vacuum filtered through a 0.2 μM PFTE membrane (Omnipore). After expression, cells were resuspended in 30 mL of 50 mM Tris-Base (pH 6.88). The resuspended cells were placed in a 10 mL falcon tube and lysed using a FB505 sonicator (1s on/off for three cycles, each 1 minute). The lysate was saturated with CO, transferred to a 30 mL round bottom tube, and centrifuged at 20,000*g* for 60 minutes to separate supernatant from non-soluble cell debris. The supernatant was collected and syringe-filtered using HPX Millex Durapore filters (Millipore) to further remove debris. A HiTrap SP cation exchange (GE) column was attached to an FPLC system (Biorad) and equilibrated in 50 mM Tris-Base (pH 6.88). The lysate was passed over the column. 50 mL of 50 mM Trise-Base (pH 6.88) was run through the SP column to remove weakly bound non-target soluble products. Elution of bound ancestral Hbs was performed with 100-mL gradient of 50mM Tris-Base 1 M NaCl (pH 6.88) buffer which was run through the column from 0% to 100%. 1.5 mL fractions were captured during the gradient process, all fractions containing red eluant were put into an Amicon ultra-15 tube and concentrated by centrifugation at 4,000g to a final volume of 1 mL. For additional purification, concentrated sample was injected into a HiPrep 16/60 Sephacryl S-100 HR size exclusion chromatography (SEC) column. The column was equilibrated in phosphate buffered saline (PBS) at pH 7.4. Purified ancestral globins elute at different volumes depending on the protein’s complex stoichiometry: 48-52 for tetramers, 56-60 for dimers, and 65-67 for monomers. The purified proteins were concentrated as mentioned above and then flash frozen with liquid nitrogen.

For experiments in which two proteins were singly expressed and purified and then mixed together, expression and purification of each protein were performed as described above. The concentration of each protein was then quantified using the Hemoglobin Assay Kit (Sigma).

Proteins were then mixed together at 50 μM each for nMS. This procedure was performed in triplicate to assess technical error introduced by the quantification and mixing process.

### Protein purification by zinc affinity chromatography

Coexpressed proteins Ancα + Ancβ were purified using zinc-affinity chromatography, which was performed using a HisTrap metal affinity column (GE) on a Biorad NGC Quest. Nickle ions were stripped from the column (buffer 100 mM EDTA, 100 mM NaCl, 20 mM TRIS, pH 8.0), followed by five column volumes of water. To attach zinc to the column, 0.1 M ZnSO_4_ was passed over until conductance was stable, approximately 5 column volumes, followed by five column volumes of water. After expression, cells were resuspended in a 50 mL lysis buffer (20 mM Tris, 150 mM Nacl, 10% glycerol (v/v), 1mM BME, 0.05% Tween-20, and 1 Roche Protease EDTA-free inhibitor tablet, pH 7.40), sonicated as described above, and the lysate passed through the prepared column. To remove non-specifically bound protein, the column was washed with 50 mL of lysis buffer. Bound protein was then eluted across a gradient of imidazole concentrations (0 to 500 mM) in a total of 100 mL elution buffer (20 mM Tris, 150 mM NaCl, 500 mM imidazole, 10% glycerol, and 1 mM BME, pH 7.4). 1 mL fractions were collected. The fraction corresponding to the second peak of UV absorbance at 280 nm has a visible red color and was collected and concentrated as described above. The concentrated solution was injected into a Biorad ENrich 650 10 x 300 columns for additional purification and eluted in PBS buffer.

### Size exclusion chromatography assay

For protein concentrations from 0 to 500 μM, size exclusion chromatography was performed using a Superdex 75 increase 10/300 GL column (GE) equilibrated in PBS, then injected with 250 μL of sample using a 2 mL injection loop on an Biorad NGC Quest FPLC and monitored by absorbance at 280 nm. For proteins at concentration 1 mM, a HiPrep 16/60 Sephacryl S-100 HR was equilibrated in PBS using an AKTAprime FPLC, then injected with 1mL sample and monitored by absorbance at 280 nm.

### Native Mass Spectrometry

Protein samples were buffer exchanged into 200mM ammonium acetate using either a centrifugal buffer exchange device (Micro Bio-Spin P-6 Gel, Bio-Rad) or a dialysis device (Slide-A-Lyzer MINI Dialysis Unit, 10000 MWCO, Thermo) prior to native MS experiments. Samples were loaded into gold-coated glass capillaries made in-house and introduced to Synapt G1 HDMS instrument (Waters corporation) equipped with a 32k RF generator (30). The instrument was set to a source pressure of 5.47 mbar, capillary voltage of 1.75 kV, sampling cone voltage of 20 V, extractor cone voltage of 5.0 V, trap collision voltage of 10 V, collision gas (Argon) flow rate of 2 mL/min (2.65 x 10 -2 mbar), and T-wave settings (velocity/height) for trap, IMS and transfer of 100 ms -1 /0.2 V, 300 ms -1 /16.0 V, and 100 ms -1/10.0 V, respectively. The source temperature (70 °C) and trap bias (30 V) were optimized. Part of the native MS experiments were conducted by Thermo Scientific Exactive Plus Orbitrap with Extended Mass Range (EMR) with tuning as follow: source DC offset of 15 V, injection flatapole DC to 13 V, inter flatapole lens to 5, bent flatapole DC to 4, transfer multipole DC to 3 and C trap entrance lens to 0, trapping gas pressure to 5.0 with the CE to 10, spray voltage to 1.50 kV, capillary temperature to 100 °C, maximum inject time to 100 ms. Mass spectra were acquired with a setting of 8750 resolution, microscans set to 1 and averaging set to 100. Mass spectra were deconvoluted using Unidec (65).

### Calculating multimerization affinity of homdimers

To estimate Kd of the monomer-to-homodimer transition of singly expressed proteins, we performed nMS at variable protein concentrations (*P_tot_*). The occupancy of each oligomeric state at each concentration was calculated as the proportion of all globin subunits in that state, based on the summed areas under the corresponding peaks in the native MS spectrum. The fraction of subunits assembled into dimers (*Fd*) includes dimers and tetramers and is defined as

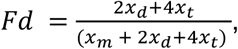

where *x_d_*, *x_t_*, and *x*_!_ are the total signal intensities of all peaks corresponding to the monomeric, dimeric and tetrameric stoichiometries, respectively. Nonlinear regression was used to find the best-fit value of Kd of dimerization using the equation:

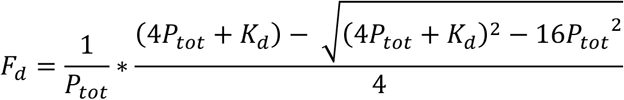

As an alternative model of homodimerization, we also used a version of the Hill-Langmuir equation:

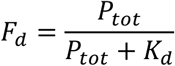

The Hill-Langmuir model, which is typically used for ligand-receptor binding, does not account for depletion of free monomeric subunits as homodimerization takes place and is therefore not a valid model in this case; we used it solely to determine the robustness of the estimated Kd to the specific binding equation used.

### Calculating multimerization affinity of homtetramers

To estimate the Kd of the homodimer– homotetramer transition, the fraction of subunits assembled into tetramers is defined as

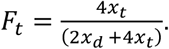

The concentration of all dimers is defined as

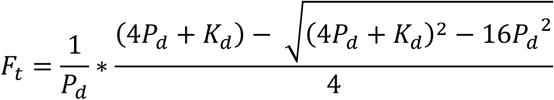

Nonlinear regression was then used to find the Kd of tetramerization using the equation:

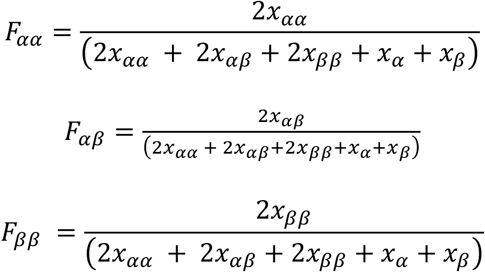

Parameters were estimated using the curvefit script in the Scipy package (66). The 95% confidence interval on the *K_d_* was estimated as 1.96 times the estimated standard error.

### Calculating multimerization affinity of heteromers

To determine the Kd of heterodimerization, we used nMS to measure stoichiometries across a titration series in which one protein’s concentration was held constant at 50 μM and the other was added at variable concentration (1 to 50 μM). From the nMS spectrum, we estimated the proportion of the heterodimer and the two homodimers as

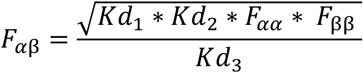

where each *x* represents the signal intensity of all peaks corresponding to the species denoted in the subscript. The dissociation constant for each dimer is defined as 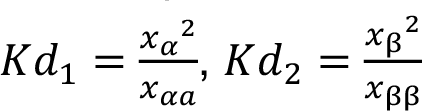, and 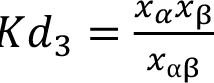. By substitution, *F_αβ_* can be expressed as

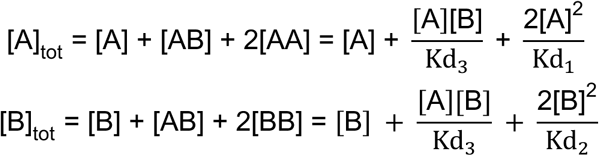

*kd*_3_ was estimated using this equation by nonlinear regression, where *F*_,,_, *F*_,2_ and *F*_22_ were measured using the titration series, and the affinities *Kd*_1_ and *Kd*_2_ were assigned the values estimated in the homodimerization experiments described above.

### Prediction of homodimer and heterodimer occupancy at high concentrations

The occupancy of each dimer at physiologically relevant concentrations (1 mM total globin subunits) was predicted as follows, because nMS is limited to concentrations <100μM. In a mixture of two types of globins *A* and *B*, the total concentration of each subunit can be expressed in terms of the concentration of monomers [A] and [B] in the mixture:

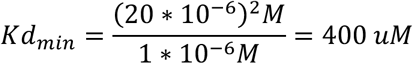

We used these equations to predict [A] and [B] at any value of C_A_ and C_B_ given the experimentally estimated Kds. The concentration of each dimer was then estimated using the equations 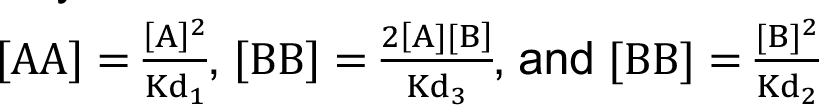.

### Establishing the upper limit of IF2 Kd

We estimated the minimum Kd of assembly across IF2 by Ancɑβ _37V+40W; IF1 removed_, because no homotetramer was observed using nMS at a protein concentration of 20 μM. The minimum detection limit for dimers in the nMS assay is 1 μM. Kd is defined as 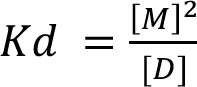, where [*M*] and [*D*] are the equilibrium concentrations of monomer and dimer, respectively. Therefore

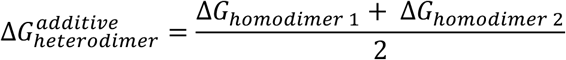

### Determining ΔΔG of specificity

Specificity for heterodimer assembly between two paralogs can be defined as the difference between the additive affinity of the heterodimer and the measured affinity of the heterodimer, using ΔGs derived measured dimerization affinity for two homodimers and their respective heterodimer. The additive affinity of the heterodimer is defined as the averaged ΔG of both homodimers:

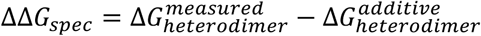

Specificity is then the difference between the additive and measured heterodimer ΔG.

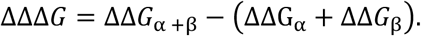

This metric is analogous to the coupling energy, which expresses the deviation of the measured DG for a double mutant from that expected given the DGs of two single mutants assuming additivity (67–69).

### Quantifying non-additive effect on specificity between Ancα and Ancβ

The non-additive effect on specificity can be defined as the difference between the predict and measured ΔΔG of the derived complex Ancα + Ancβ.

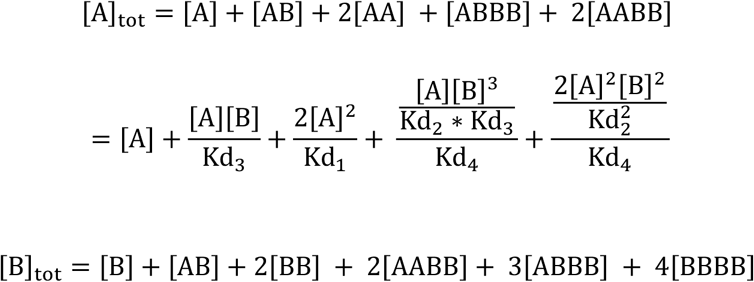

### Prediction of monomer, dimer, and tetramer occupancies with no IF2 specificity

The occupancy of monomers, dimers, and tetramers between 1 mM and 4 mM predicted was calculated as follows. The concentration of subunit in each stoichiometric species can be expressed in terms of the concentration of monomers [A] and [B]:

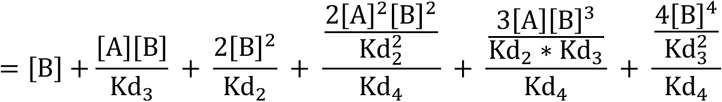

We used these equations to predict [A] and [B] across a range of [A]_IJI_ and [B]_IJI_ values given previously measured equilibrium constants. Predicted [A] and [B] concentrations were used to calculate the concentration of homodimers and heterodimers as described above, and the concentration of tetramers were calculated using the following equations:

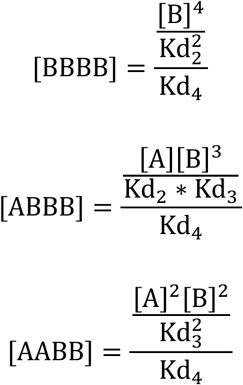

where [BBBB] corresponds to the concentration of homotetramer, [ABBB] is concentration of ⍺_0_β_1_ tetramers, and [AABB] is the concentration of ⍺_#_β_#_ heterotetramers.

### Homology models

SWISS-Model was used to generate a structural model of the Ancαβ_q40W_ homotetramer using the crystal structure of the human Hbβ homotetramer (PDB 1CMB) as template, which was then refined using Rosetta’s Fast Relax protocol, which energetically minimizes the initial structure via small adjustments to the backbone and side chain torsion angles (70). PyMOL V2.1 was used to visualize the proteins and capture images.

IF1-mediated homodimers were generated by the same procedure, except for homodimers of Ancα or Ancαβ_ΔD_, for which the homodimer of human Hbα (PDB 3S48) was used as template. IF1-mediated heterodimers were generated by the same procedure but using the heterotetramer of human Hb (PDB 4HHB). For PyMol PSE file containing these models, see Supplementary file.

## Acknowledgements and funding sources

We thank members of the Thornton and Laganowsky labs for helpful advice and comments. Supported by NIGMS R01GM131128, R35 GM14533601, 1T32GM13978201.

## Author Contributions

CRCR: project conception and design, protein expression and purification, chromatography experiments, structural modeling, data analysis, and writing (initial draft and revisions). JL: design, execution, and analysis of mass spectrometry experiments, and writing (revisions). AP: project conception and design, protein expression and purification, chromatography experiments, data analysis, writing (initial draft and revisions). AL: design and analysis of mass spectrometry experiments, writing (revisions). JWT: project conception and design, data analysis, project management, funding acquisition, writing (initial draft and revisions).

## SUPPLEMENTARY FIGURES

**Fig. S1.**
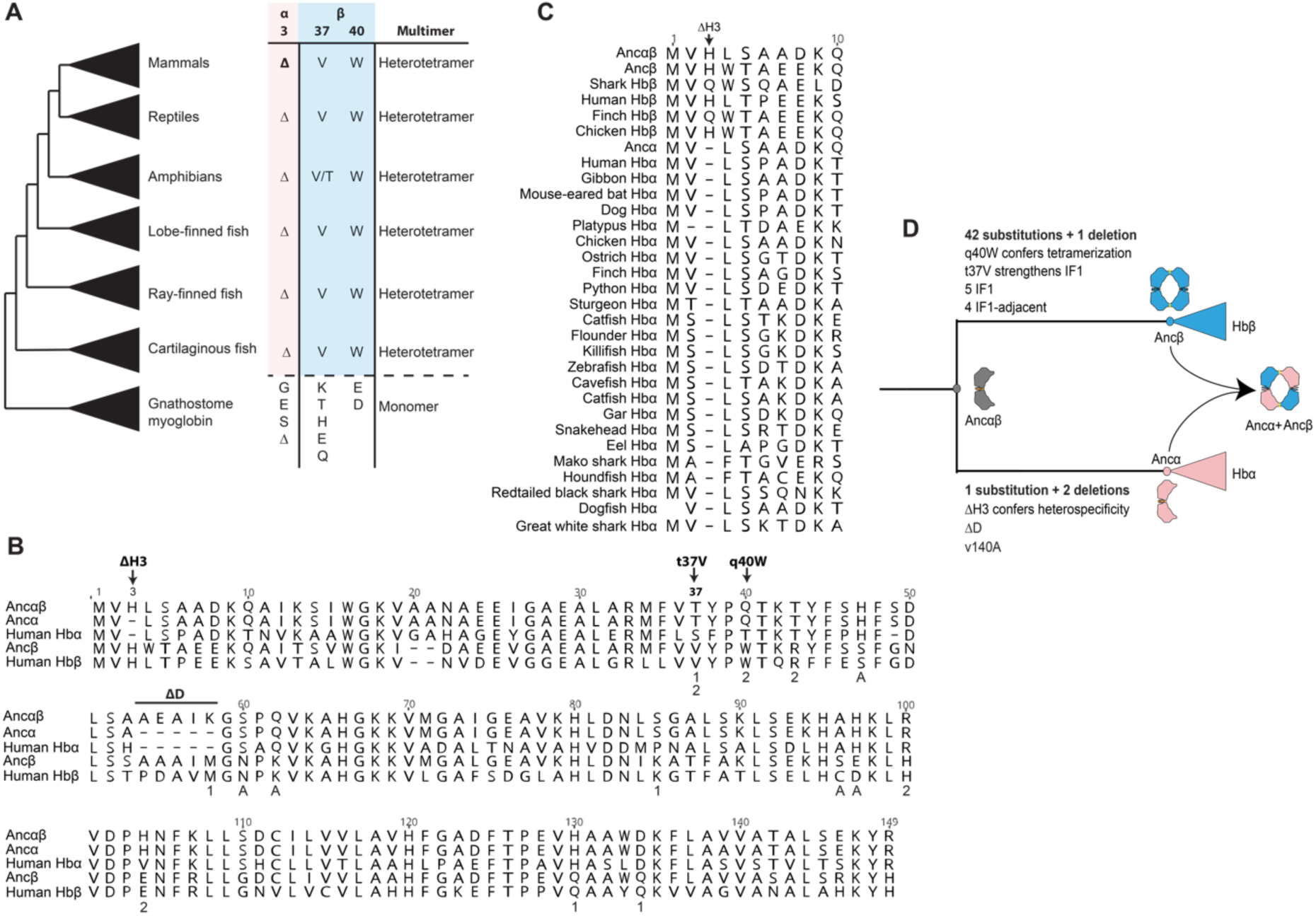
Key sequence changes are conserved in extant hemoglobin subunits. **A**) Multimerization state and residues at key sequence sites in extant Hb subunits from major jawed vertebrate taxa. Myoglobin is the closest paralogous protein in the globin family. For complete alignment, see https://tinyurl.com/yc6kvdb8. **B**) Alignment of extant human and reconstructed ancestral Hb subunits is shown. Historical sequence changes that confer heterospecificity (Λ1H3 in Hbα) and tetramerization (q40W in Hbβ) are labeled and shown with arrows. Additional substitutions that occurred on the branch leading to Ancβ and assayed in this paper are also labeled, including changes on the surface of IF1 (1), IF2 (2), and at sites adjacent to IF1 (A). **C)** Alignment of N-terminal portion of Hb subunits from species representative of major vertebrate taxa. The deletion of residue 3 in alpha subunits is marked. **D**) Summary of key sequence changes that occurred after the duplication of Ancαβ. Multimerization states of reconstructed ancestral proteins is shown.

**Fig. S2.**
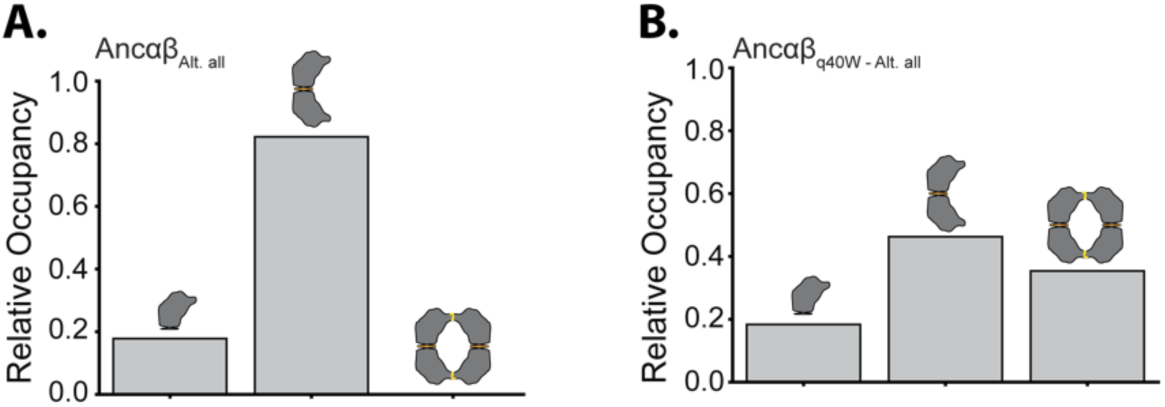
Effect of q40W tetramerization is robust to statistical uncertainty. (A) Relative occupancy of monomer, dimer, and tetramer of Ancɑβ_Alt. all_, an alternative reconstruction of Ancɑβ that contains the second most likely state at all ambiguously reconstructed sites, measured at 20 µM total protein using native MS. (B) Relative occupancy Ancɑβ_Alt. all_ with substitution q40W.

**Fig. S3.**
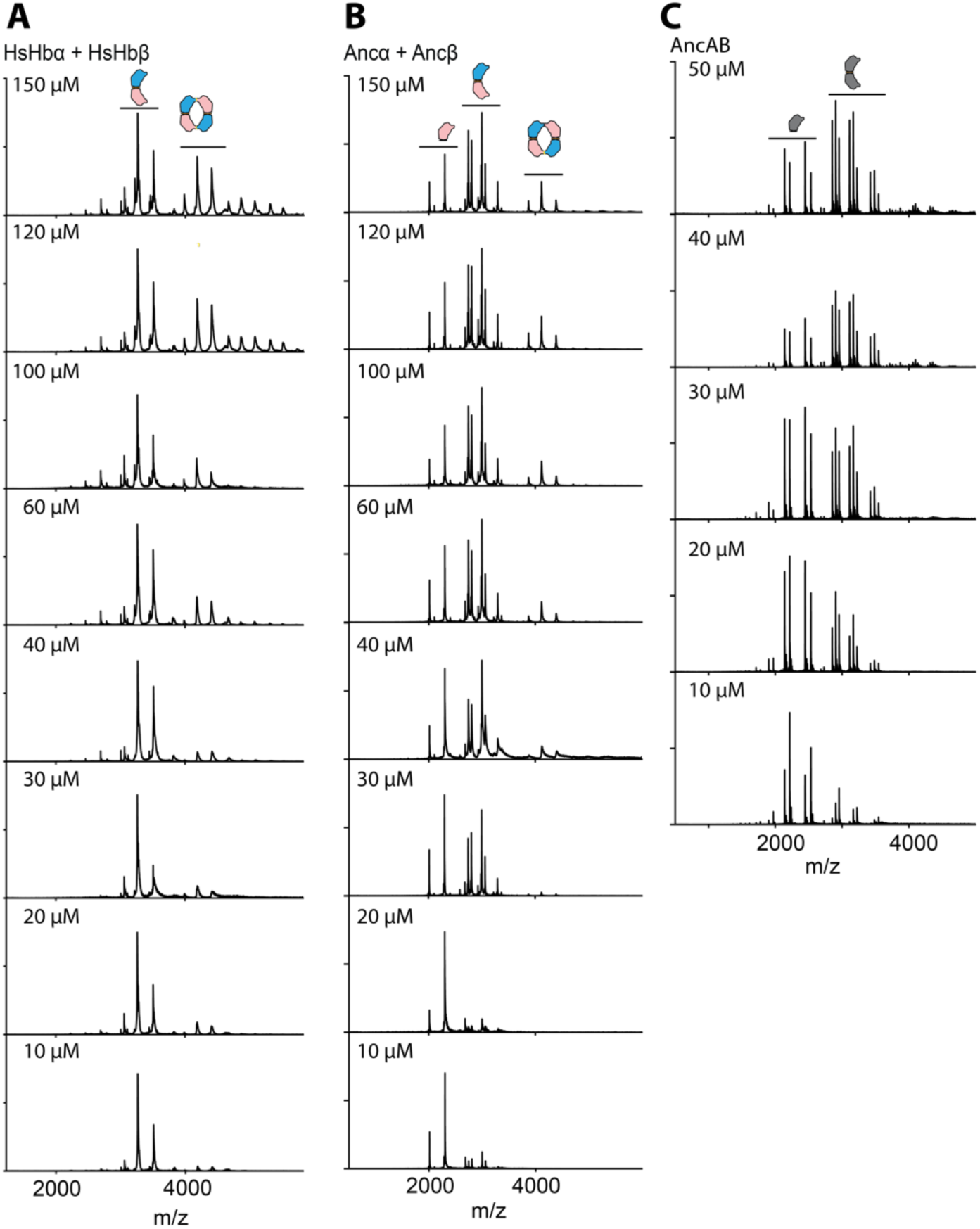
Native mass spectrometry spectra. nMS spectra across a concentration series is shown for A) human Hb, B) Ancɑ + Ancβ, and (C) Ancɑβ. Peaks corresponding to monomers, dimers, and tetramer are labeled.

**Fig. S4.**
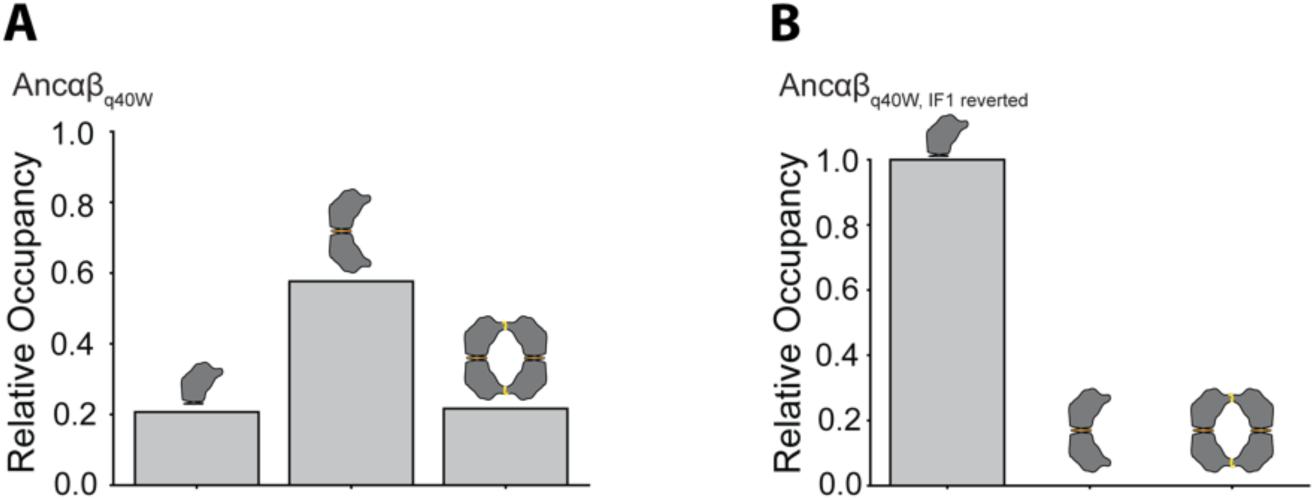
The effect of q40W on tetramerization depends on IF1. (A) Relative occupancy of Ancɑβ_q40W_, measured by native MS at 20 µM total protein. (B) Relative occupancy of Ancɑβ_q40W_IF1-reverted_, which contains mutation q40W, as well as reversions to the ancestral state found in AncMH of all residues that were substituted between AncMH and Ancαβ.

**Fig. S5.**
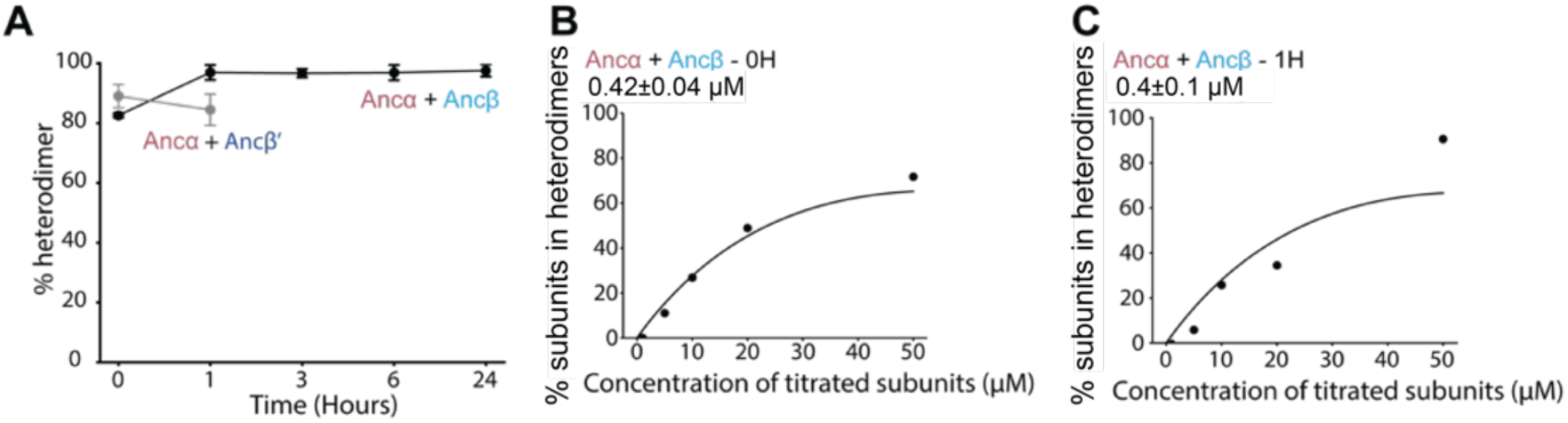
Heterodimer occupancy of Ancα and Ancβ is near equilibrium after mixing. (A) The percent of all dimers that are heterodimers, measured by nMS when proteins are mixed at 50 µM each and allowed to incubate for 0, 1, 3, 6, or 24 hours. Black line and points, Ancα + Ancβ (which only dimerize when expressed separately and then mixed). Grey line and points, Ancα + Ancβ’ (Ancβ in which IF2 surface substitutions are reverted to their ancestral state in Ancαβ, thus preventing tetramerization). Each dot shows the mean of three replicates; error bars, SEM. (B) Affinity of monomer-to-heterodimer assembly measured by nMS immediately upon mixing of Ancα and Ancβ. Ancα was kept constant at 50 µM, while the concentration of Ancβ varied. Points, fraction of all subunits in the mixture that are incorporated into heterodimers. Line, best-fit binding curve. Estimated Kd and 95% confidence interval are shown. (C) Estimated heterodimerization affinity measured as in panel B, but 1 hour after mixing.

**Fig. S6.**
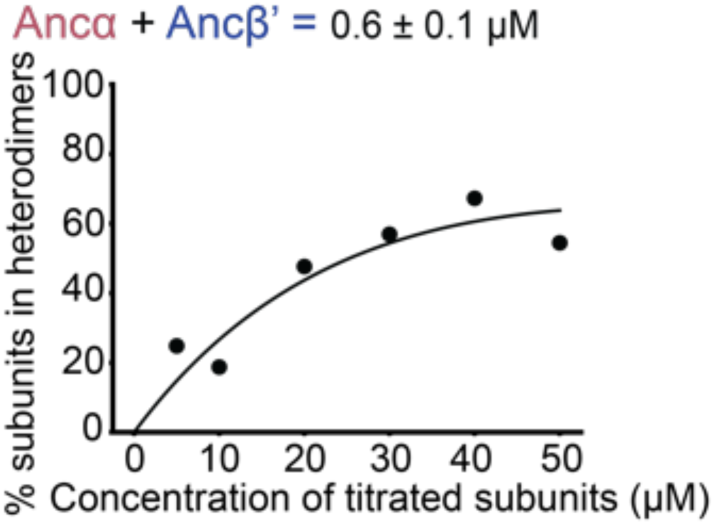
Heterodimerization by Ancα+Ancβ’. Monomer-to-heterodimer assembly measured by nMS. Ancα was kept constant at 50 µM while Ancβ’ was at variable concentration. Points, fraction of all subunits in the mixture that are incorporated into heterodimers at each concentration. Line, best-fit binding curve. Estimated Kd and 95% confidence interval are shown.

**Fig. S7.**
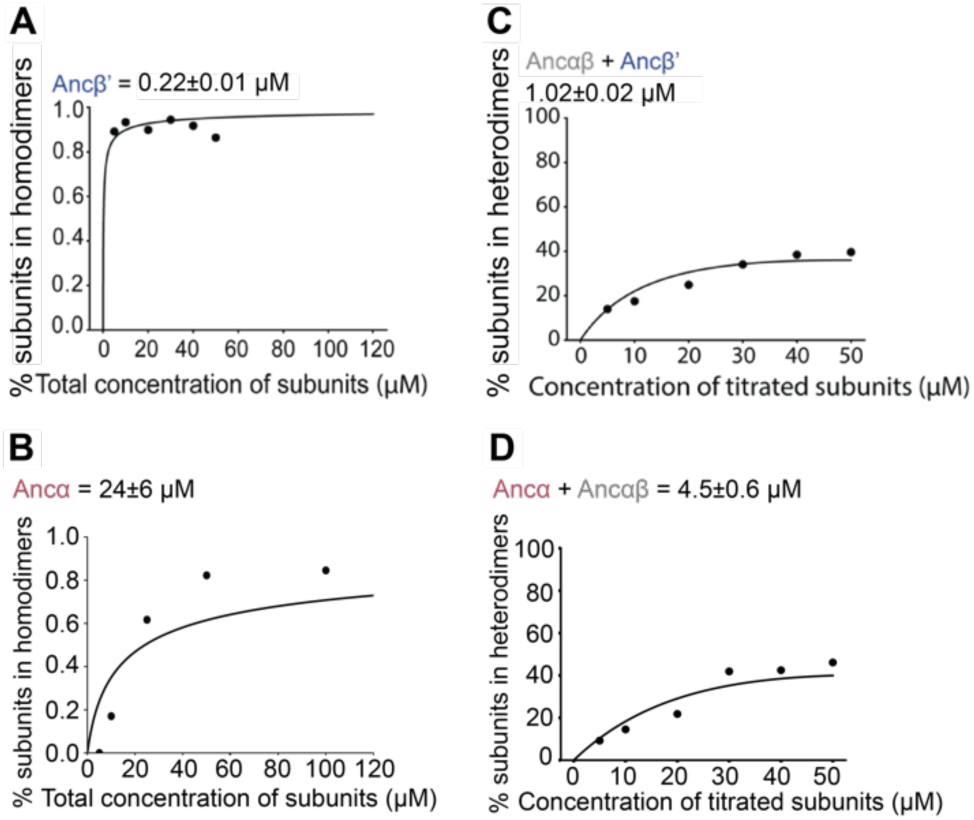
Dimerization by Ancα and Ancβ’. (A-B) Homodimerization by Ancβ’ (panel A) and by Ancα (B). measured by nMS across a titration series. Each point shows the fraction of subunits incorporated into dimers as the concentration of protein varied. Best-fit binding curve, Kd, and 95% confidence interval are shown. (C-D) Heterodimerization by mixtures of Ancαβ+ Ancβ (C) and Ancαβ+Ancα and Ancα+Ancαβ (D). Each point shows the fraction of all subunits incorporated into heterodimers. In each case, one protein was held constant at 50 μM while the other was varied.

**Fig. S8.**
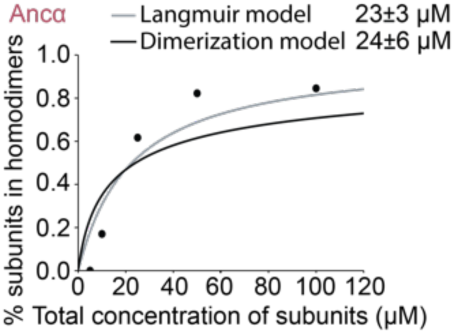
Estimated affinity of the monomer-dimer transition by the Ancα homodimer is robust to the binding model used. We fit two different binding equations (curves) to the stoichiometries of **Ancα** measured using nMS. The Langmuir-Hill model (black line) assumes that the concentration of free monomeric subunits is not depleted by dimerization. The dimerization model used throughout the rest of this paper accounts for this depletion (described in Materials and Methods). The estimated Kds (shown with their 95% confidence intervals) are statistically indistinguishable.

**Fig. S9.**
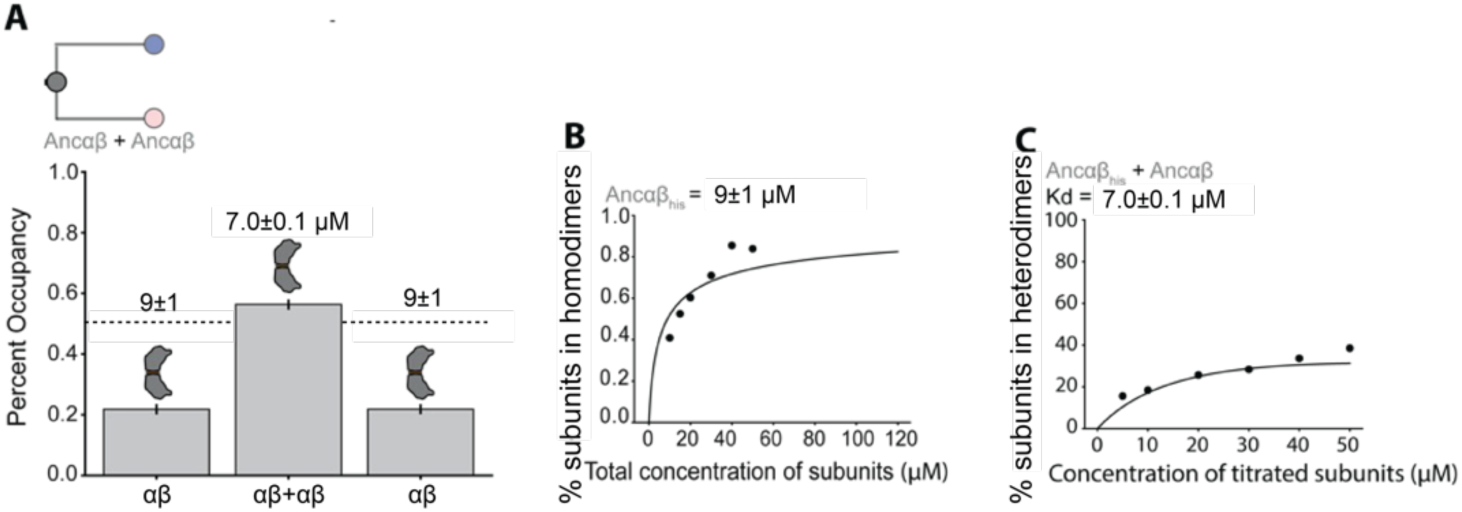
Dimerization affinities and occupancy of Ancαβ. (A) Expected fractional occupancies of homodimer and heterodimers when Ancαβ is mixed at equal concentrations with Ancαβ_his_ (500 μM each), given the measured dimerization affinities (shown above each column, with 95% confidence interval). Ancαβ_his_ is Ancαβ with an N-terminal polyhistidine tag, which allows the masses of the three kinds of dimer to be distinguished. (B-C) Homodimerization by Ancαβ_his_ and heterodimerization by affinity of Ancαβ+ Ancαβ_his_, measured and represented as in Fig. S5.

**Fig. S10.**
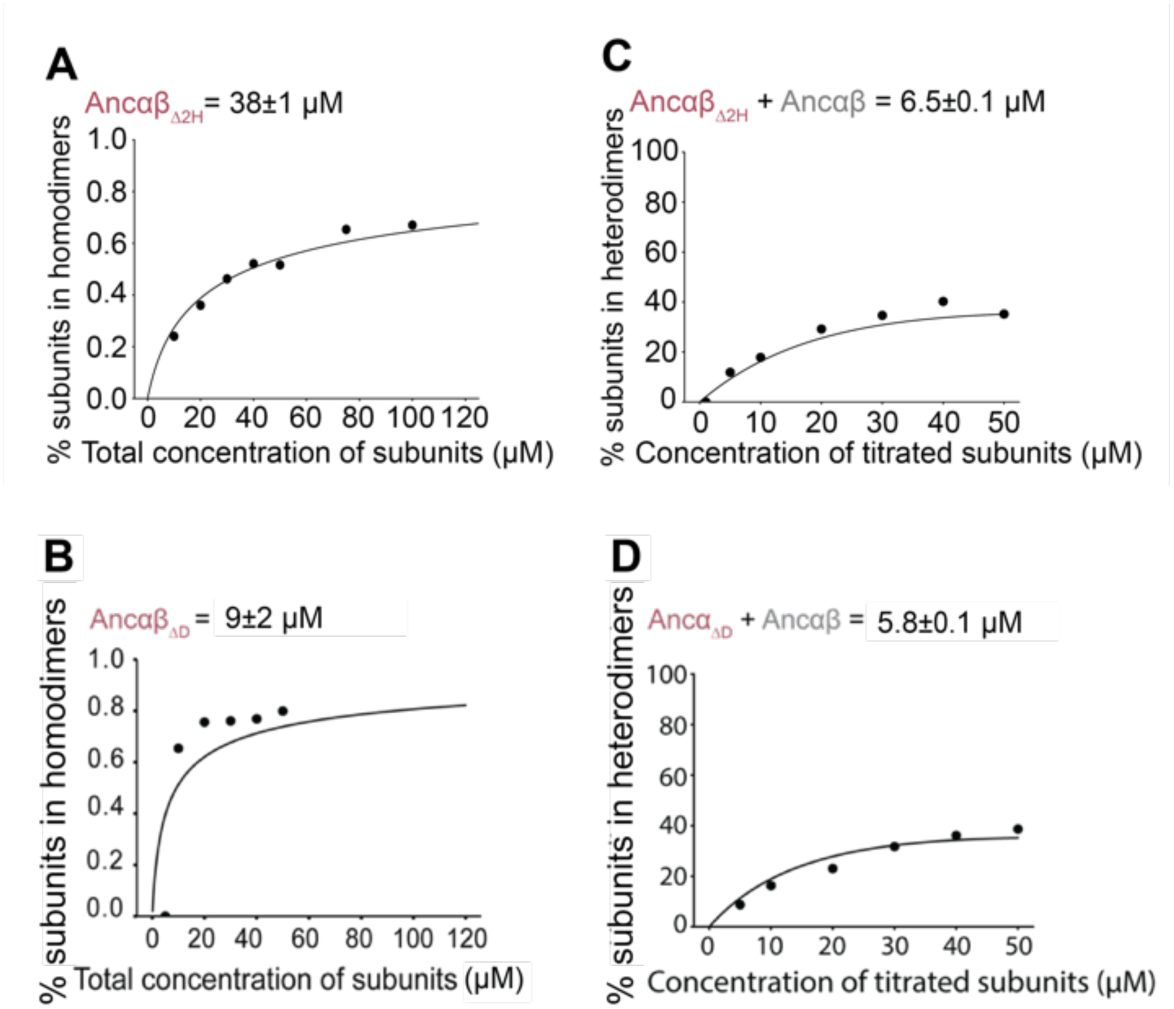
Effect of historical deletions on dimerization. (A-B) Homodimerization and (C-D) Heterodimerization by mixtures, measured and represented as in Fig. S5.

**Fig. S11.**
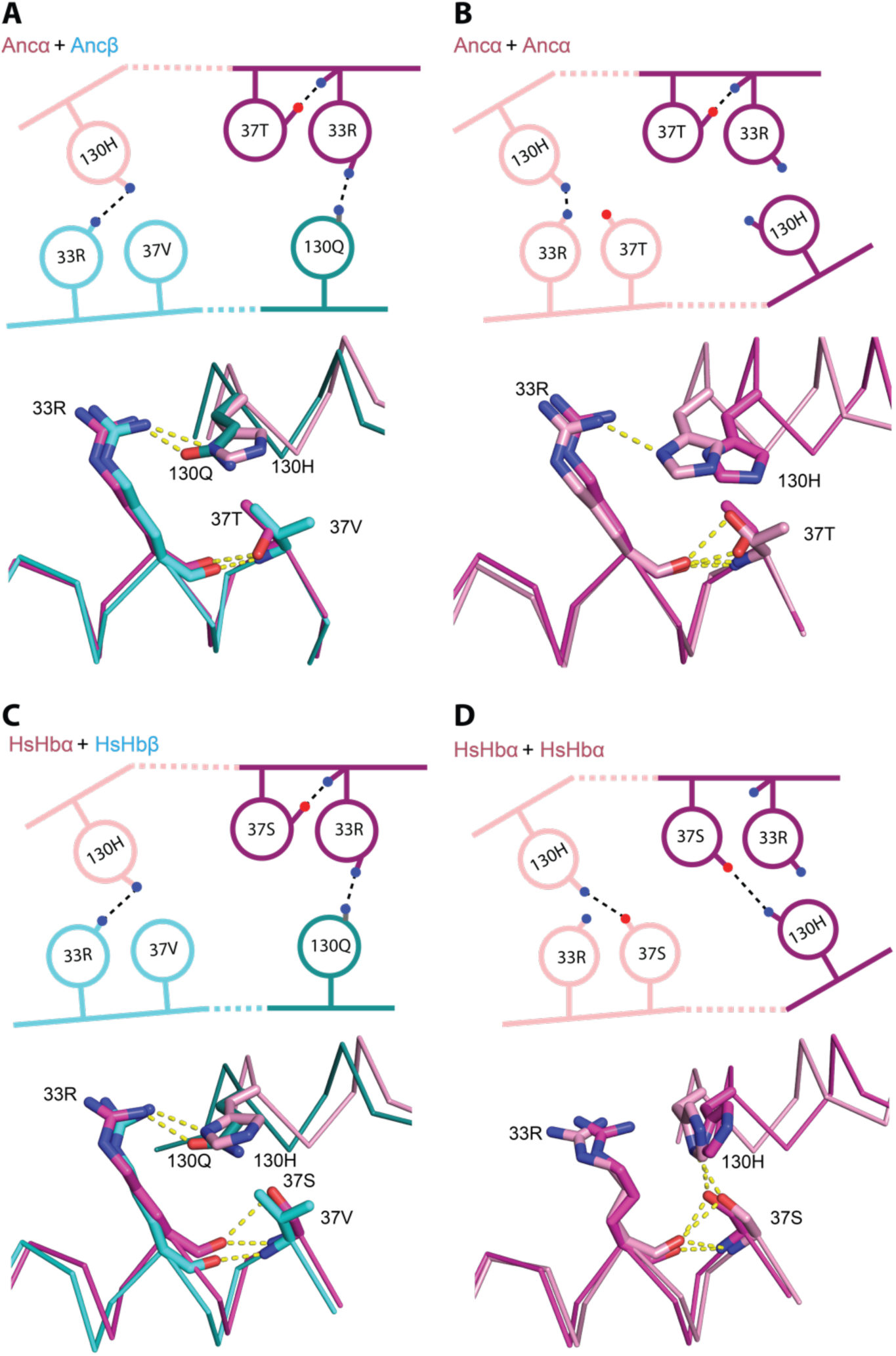
Nonadditive interactions that contribute to specificity are conserved in derived Hb complexes. In the modeled homodimers and heterodimers of Ancɑ+Ancβ (panels A, B) and X-ray crystal structure of human hemoglobin (PDB 4HHB and 3S48), the figure shows the key IF1 residues with nonadditive interactions in Ancɑβ+AncɑβΔH3 (see Fig. 5G for comparison). Top, cartoon of key contacts. The two iterations of these interactions across the isologous interface are shown, one each in light or dark hue. Blue and red, nitrogen and oxygen atoms, respectively. Dotted lines, hydrogen bonds. Bottom, structural alignment of the two iterations of the isologous interface in each dimer. Each dimer structure was duplicated exactly and then aligned to the original by targeting one subunit of the copy to align to the other subunit of the original. Hues correspond to the isologous iterations in the cartoon above

**Fig. S12.**
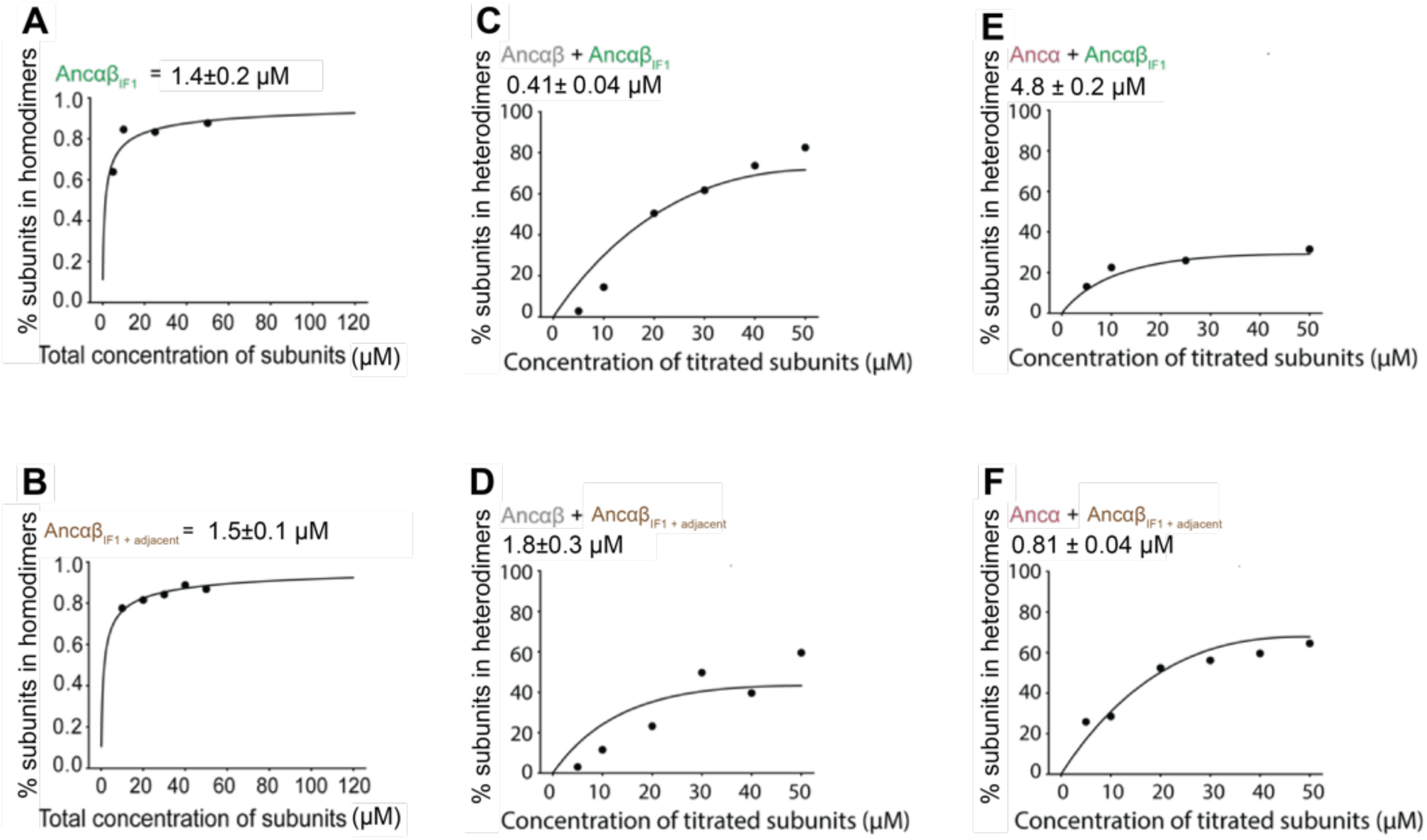
Homodimerization by Ancαβ_IF1_ and Ancαβ_IF1 + Adjacent_. (A,B) and heterodimerization by those proteins when mixed with Ancαβ (C,D) or Ancα (E,F). Measurements and representation as in Fig. S5.

**Fig. S13.**
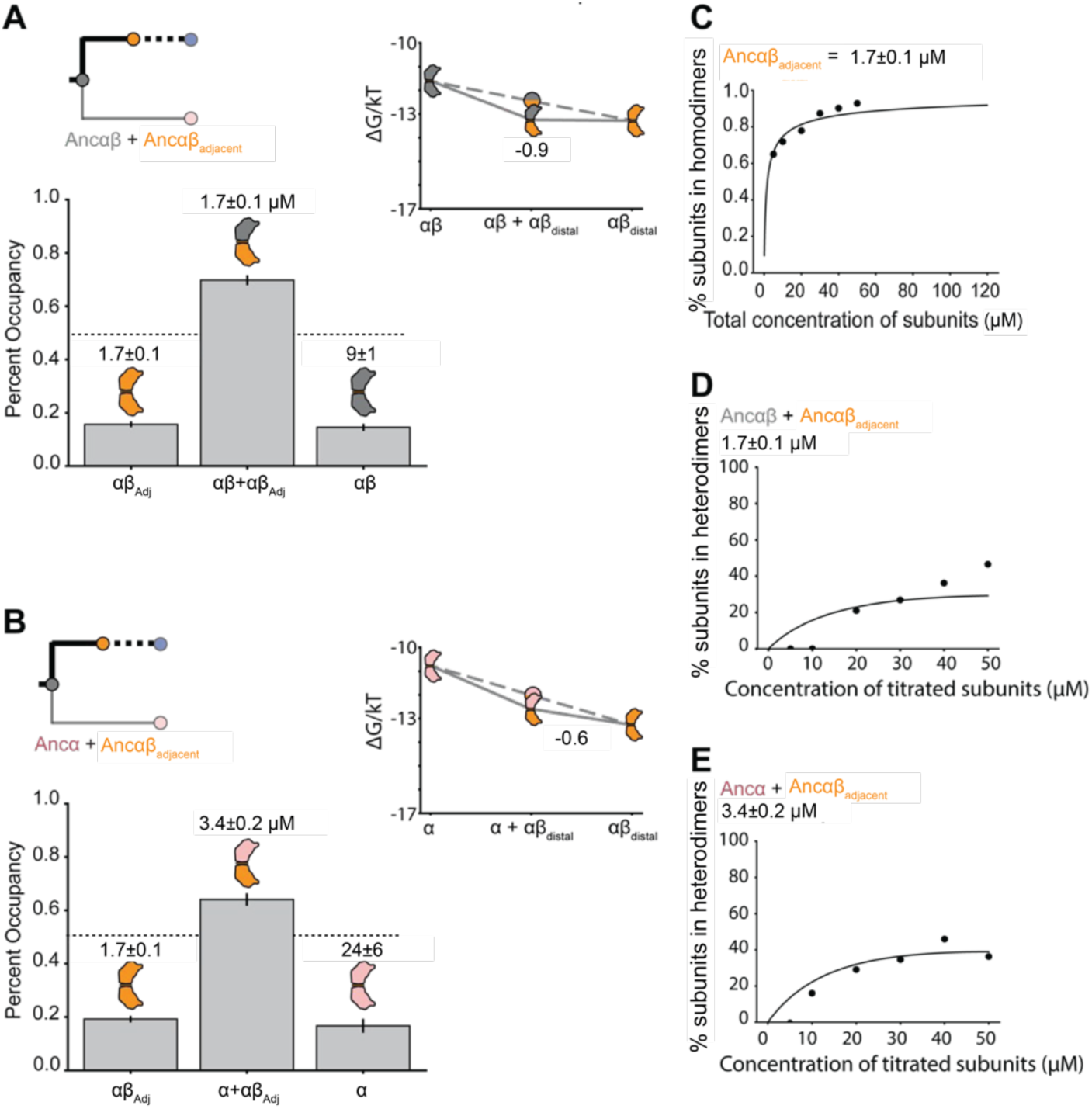
Dimerization affinity and occupancies for Ancαβ_Adjacent_. Expected fractional occupancies of homodimer and heterodimers when Ancαβ_Adjacent_ Is mixed with Ancαβ (A) or Ancα (B), each at (500 μM), given the measured dimerization affinities (shown above each column, with 95% confidence interval). Inset, ΔG of each dimerization (measured in units of kT), with ΔG_spec_ of the heterodimer shown. (C,D,E) Measurement of binding affinities, measured and represented as in Fig. S5.

